# NK cell-monocyte crosstalk underlies NK cell activation in severe COVID-19

**DOI:** 10.1101/2023.10.27.564440

**Authors:** MJ Lee, I de los Rios Kobara, TR Barnard, X Vales Torres, NH Tobin, KG Ferbas, AW Rimoin, OO Yang, GM Aldrovandi, AJ Wilk, JA Fulcher, CA Blish

## Abstract

NK cells in the peripheral blood of severe COVID-19 patients exhibit a unique profile characterized by activation and dysfunction. Previous studies have identified soluble factors, including type I interferon and TGFβ, that underlie this dysregulation. However, the role of cell-cell interactions in mediating changes in NK cell function during COVID-19 remains unclear. To address this question, we combined cell-cell communication analysis on existing single-cell RNA sequencing data with *in vitro* primary cell co-culture experiments to dissect the mechanisms underlying NK cell dysfunction in COVID-19. We found that NK cells are predicted to interact most strongly with monocytes and that this occurs via both soluble factors and direct interactions. To validate these findings, we performed in vitro co-cultures in which NK cells from healthy donors were incubated with monocytes from COVID-19+ or healthy donors. Co-culture of healthy NK cells with monocytes from COVID-19 patients recapitulated aspects of the NK cell phenotype observed in severe COVID-19, including decreased expression of NKG2D, increased expression of activation markers, and increased proliferation. When these experiments were performed in a transwell setting, we found that only CD56^bright^ CD16^-^ NK cells were activated in the presence of severe COVID-19 patient monocytes. O-link analysis of supernatants from transwell co-cultures revealed that cultures containing severe COVID-19 patient monocytes had significantly elevated levels of proinflammatory cytokines and chemokines as well as TGFβ. Collectively, these results demonstrate that interactions between NK cells and monocytes in the peripheral blood of COVID-19 patients contribute to NK cell activation and dysfunction in severe COVID-19.

**BACKGROUND:** Natural killer (NK) cells are innate lymphocytes that are critical antiviral effectors. Because of their role in controlling acute viral infections, multiple studies have evaluated the role of NK cells in SARS-CoV-2 infection. Such studies revealed that NK cell phenotype and function are significantly altered by severe COVID-19; the peripheral NK cells of severe COVID-19 patients are highly activated and proliferative(1–5), with increased expression of cytotoxic molecules, Ki-67, and several surface markers of activation(3, 5–8). However, these NK cells also have dysfunctional cytotoxic responses to both tumor target cells(1, 2, 9, 10) and SARS-CoV-2-infected target cells(9, 10). Given that peripheral NK cells are thought to migrate to the lung during COVID-19(11–13), these results suggest that the NK cells of severe COVID-19 patients may be incapable of mounting a successful antiviral response to SARS-CoV-2 infection.

Although the unique phenotype and dysfunctionality of NK cells in severe COVID-19 has been well-characterized, the processes underlying these phenomena have not. Only one study has conducted *in vitro* mechanistic experiments to identify a possible cause of NK cell dysfunction: Witkowski et al. identified serum-derived TGFβ as a suppressor of NK cell functionality in severe COVID-19 patients(9). However, this study did not identify the source of serum TGFβ. Additionally, given the high degree of complexity within the immune system, there are likely other causes of NK cell dysfunction in COVID-19 that have thus far remain unexplored. One such mechanism may be the myriad of interactions between NK cells and other peripheral immune cells. NK cells are known to interact with CD4 and CD8 T cells, dendritic cells, neutrophils, and macrophages/monocytes(14), which can prime NK cell cytotoxicity or induce tolerance. Previous work by our lab suggested the potential for NK cell-monocyte crosstalk in severe COVID-19 through the expression of ligands for NK cell activating receptors on the monocytes of these patients(3). Crosstalk between NK cells and monocytes plays a role in regulating the NK cell response to other infections, including HIV-1(15, 16), mouse(17) and human cytomegalovirus(18), and malaria(19) through mechanisms including secretion of NK cell-regulating cytokines by monocytes.

In this study, we used a combination of computational and *in vitro* methods to dissect the interactions between NK cells and monocytes in severe COVID-19. We utilized primary NK cells and monocytes from a large cohort of COVID-19 patients to demonstrate that co-culture of healthy NK cells with monocytes from severe COVID-19 donors can partially recapitulate the activated phenotype observed in the NK cells from COVID-19 patients. We then interrogated the mechanisms by which this activation occurs by performing NK cell-monocyte co-cultures in a transwell setting and using O-link to analyze the cytokines present in this system. Collectively, our work identifies monocytes as a driver of NK cell activation in severe COVID-19 and reveals interactions between NK cells and monocytes that may underlie this process.

## RESULTS

### Transcriptomic analysis reveals NK cell-monocyte crosstalk in severe COVID-19

We began by identifying the cell types interacting with NK cells in severe COVID-19. To do so, we first probed the cell-cell communication pathways that may contribute to the NK cell activation, proliferation, and dysfunction in severe COVID-19 using <u>S</u>ingle-<u>C</u>ell <u>R</u>esolution <u>A</u>nalysis through <u>B</u>inning (“Scriabin”)(20). We applied this algorithm to a previously acquired COVID-19 dataset(3) to identify the interactions in which the NK cells were the receiving cell type and the cell types that were the predicted senders of these signals. To do this, we applied Scriabin’s interaction program (IP) discovery workflow, which identifies groups of ligands and receptors that are significantly co-expressed by the same sets of sender and receiver cells. These IPs thus represent modules of highly topologically connected cell-cell communication pathways. When generating IPs with NK cells acting as the receiver cell type, we found that monocytes had the highest co-expression of ligands (both soluble and membrane-bound) for NK cell receptors (Fig. 1A). Interestingly, even in healthy donors, NK cells received most signals from monocytes; however, the magnitude of these interactions was much greater in the setting of severe COVID-19. We therefore focused our efforts on analyzing interactions between NK cells and monocytes.

**Fig. 1:**
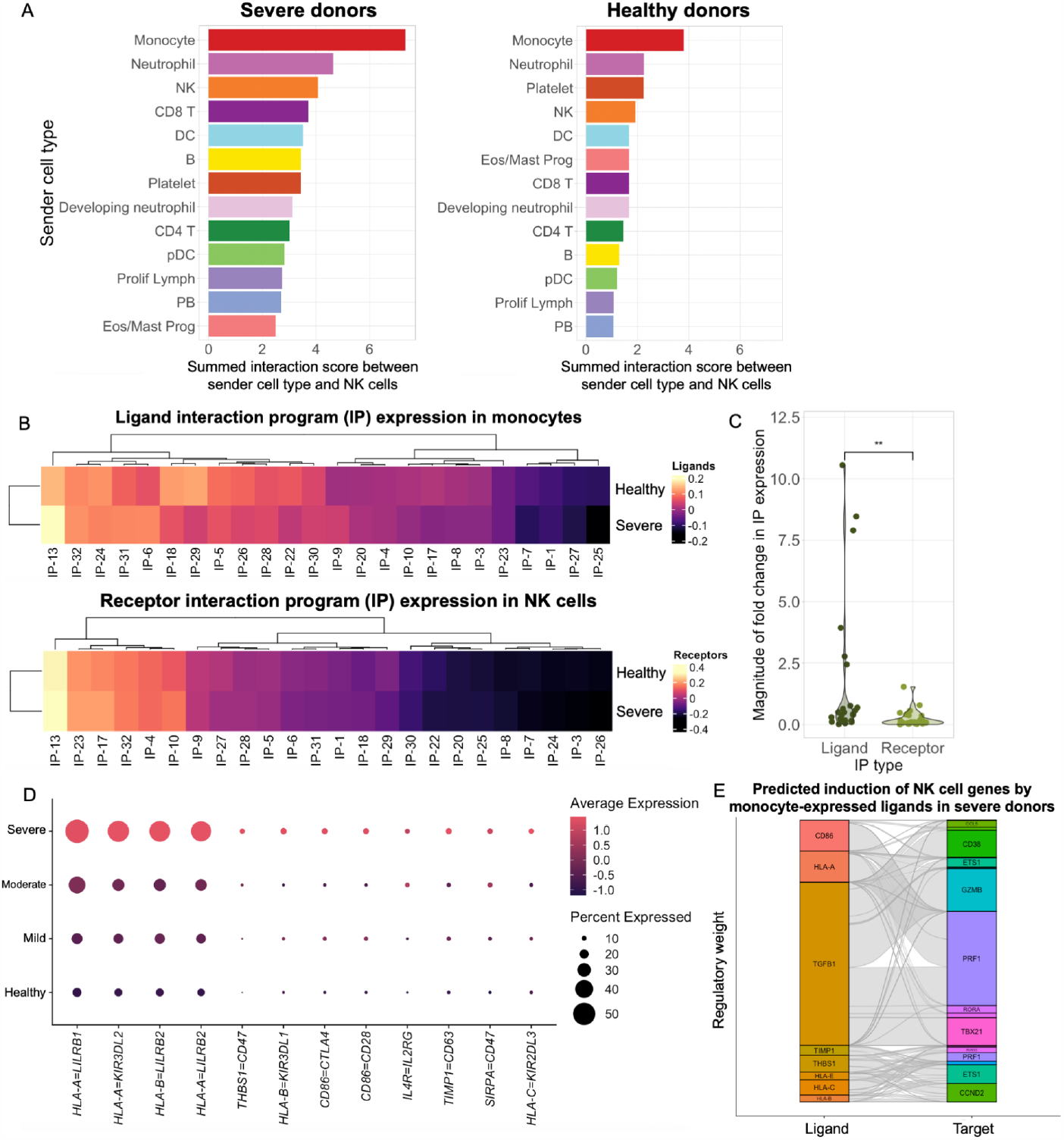
Robust monocyte-NK cell crosstalk in severe COVID-19. A scRNA-seq dataset from Wilk & Lee et al. 2021 was analyzed using the R package *Scriabin*. A) Bar plots showing the summed interaction score between a given sender cell type (shown on y-axis) and NK cells. A larger score indicates more individual points of interaction between the sender cell type and NK cells. Interactions between cell types are shown separately samples from severe COVID-19 donors (left) and healthy control donors (right). B) Heatmaps showing the scaled expression of each interaction program (IP) that is significantly expressed in NK cells and monocytes from severe COVID-19 and healthy donors. Top heatmap shows scaled expression of the ligands for each interaction program in monocytes. Bottom heatmap shows scaled expression of the receptors for each interaction program in NK cells. C) Violin plot showing the magnitude of fold change in expression for each IP in monocytes (“Ligand”) and NK cells (“Receptor”) between healthy and severe COVID-19 samples. Magnitudes of fold change were calculated by taking the absolute value of the fold change in expression of each IP between healthy and severe samples. Significance value was calculated using a Wilcoxon signed-rank test. D-E) Dotplots showing weighted expression of differentially-expressed ligand-receptor pairs in monocytes (ligand expression) and NK cells (receptor expression) across COVID-19 severities. Ligand-receptor expression was weighted by predicted activity using the *NicheNet* package(21). F) Alluvial plot showing predicted expression of genes in NK cells (right) based on weighted expression of ligand-receptor pairs from subfigure D on monocytes and NK cells (left). Larger ligand boxes are predicted to have more regulatory weight in driving downstream signaling. Larger target boxes are predicted to be more strongly regulated by the active ligands. Gray lines illustrate relationships between regulatory ligands and downstream targets.

Given the strong monocyte-NK cell crosstalk suggested by this analysis, we next explored what ligand-receptor pairs underpinned these interactions, and their predicted effects on downstream NK cell transcriptional phenotype. We performed this analysis on NK cells and monocytes from healthy and severe COVID-19 donors and found 24 IPs that were significantly co-expressed by either healthy donor cells or severe COVID-19 donor cells (Table 1). Heatmaps showing the scaled expression of the ligands for these programs in monocytes or the receptors for these programs in NK cells demonstrate that the bulk of the changes in severe COVID-19 patients occur in monocyte expression of ligands rather than in NK cell expression of receptors (Fig. 1B). Quantifying the magnitude of fold change in each IP between healthy and severe donors also shows that there are significantly greater changes in ligand IP expression than in receptor IP expression (Fig. 1C).

We determined which ligand-receptor pairs were more active in severe COVID-19 patients compared to healthy controls using Scriabin’s Cell-Cell Interaction Matrix (CCIM) workflow, which creates a matrix of ligand-receptor pair expression for each pair of individual cells within the dataset. We found 12 edges whose weighted expression was increased in the NK cells and monocytes of severe COVID-19 patients (Fig. 1D). Interestingly, half of these edges consisted of a classical MHC class I ligand and an inhibitory NK cell receptor (Fig. 1D). Weighted expression of the costimulatory molecule CD86 and its cognate receptors, immune checkpoint molecule CTLA-4, and activating receptor CD28, were also increased in severe patients (Fig. 1D).

Finally, we used Scriabin to identify NK cell genes predicted to be upregulated by ligands that are highly expressed on the monocytes of severe COVID-19 patients. We analyzed the predicted downstream targets of 6 ligand-receptor pairs that were significantly upregulated in severe COVID-19 patients compared to healthy controls (HLA-A/B/C, CD86, TIMP1, and THBS1) (Fig. 1E). We also included TGFB1, which is expressed by both NK cells and monocytes and has been shown to play a role in NK cell dysfunction in COVID-19(9). Collectively, these ligands were predicted to induce expression of genes that are known to be expressed in NK cells from severe COVID-19 patients, including *GZMB, PRF1*, and *CD38(3, 5)*. Other predicted targets include the transcription factors *ETS1* and T-bet (*TBX21*), both of which regulate NK cell function and survival (Fig. 1E), and the cyclin *CCND2* (cyclin D2) which is involved in cellular proliferation(22). Altogether, these analyses demonstrate that monocytes interact with NK cells, that these interactions are stronger during severe COVID-19, and nominate monocyte-expressed ligands such as CD86 and TGFβ as drivers of known NK cell phenotypes in COVID-19.

### NK cells from hospitalized COVID-19 patients are activated and proliferative

Having identified monocytes as a cell type that interacts strongly with NK cells in COVID-19, we next sought to devise an *in vitro* experimental system that would allow us to interrogate NK cell-monocyte interactions using primary immune cells from COVID-19-positive donors. For this, we used a cohort of 44 hospitalized donors collected from an observational cohort study at UCLA as well as 17 healthy donors from the Stanford Blood Bank (Fig. 2A-B). Given that previous studies, including by our group, have identified striking phenotypic differences in the NK cells of hospitalized COVID-19 patients compared to those of mild COVID-19 patients and healthy controls(1, 4–6, 8–10, 23, 24), we first sought to assess phenotypic changes in the NK cells of this cohort of COVID-19 patients. Similar to prior studies, we found that COVID-19 induces a shift in the frequency of NK cell subsets defined by expression of CD56 and CD16. NK cells from COVID-19 patients had a significant increase in the frequency of unconventional CD56^dim^ CD16^lo^ NK cells and a corresponding decrease in the frequencies of both CD56^dim^ CD16^hi^ and CD56^bright^ CD16^lo^ NK cells (Fig. 2C-D).

**Fig. 2:**
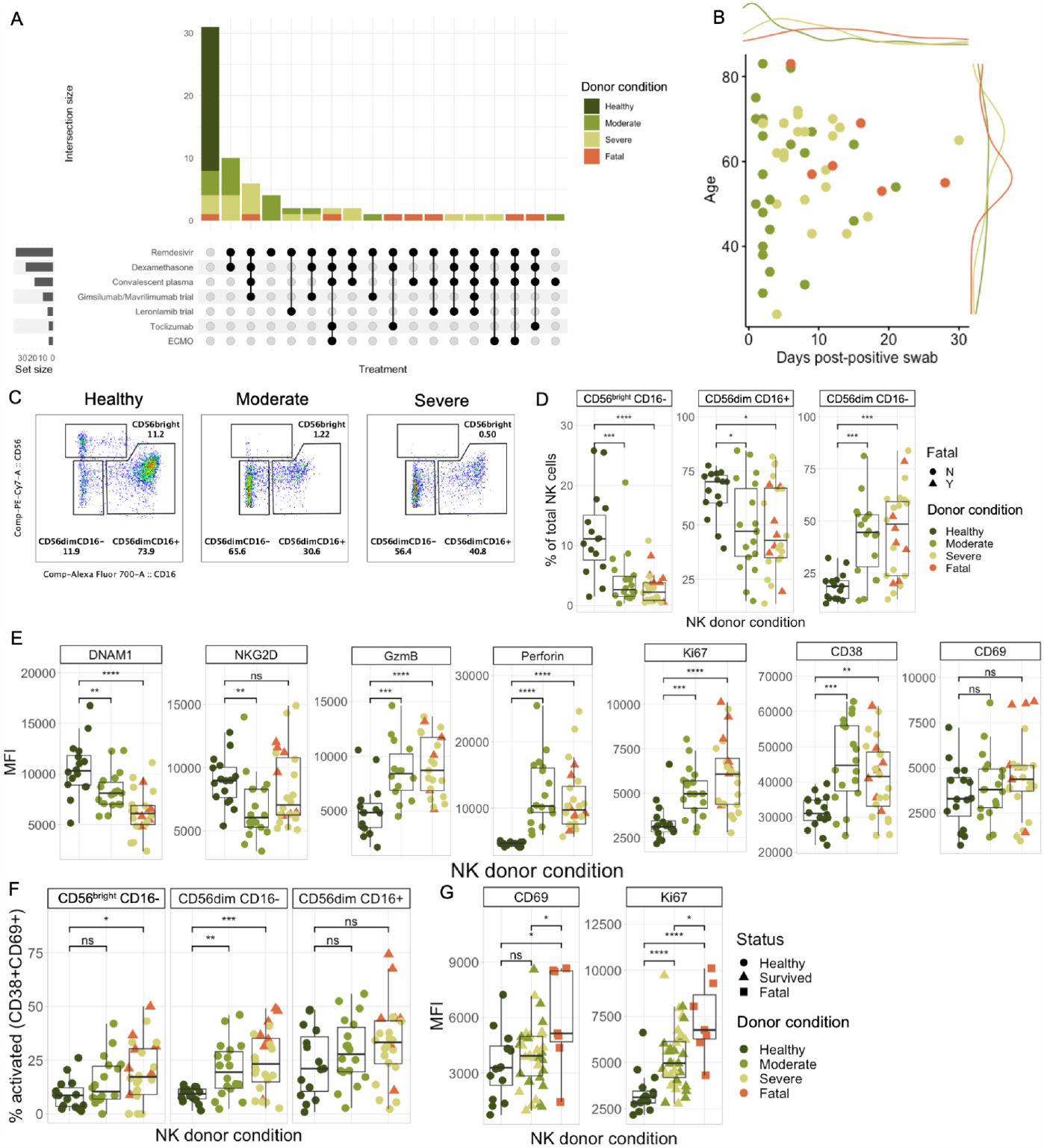
NK cells from hospitalized COVID-19 patients are phenotypically altered. A) Upset plot showing the number of patients in each treatment group, colored by patient severity. B) Scatter plot showing the distribution of age (y axis) and days post-positive test swab (x axis) within the cohort of COVID-19-positive donors. C-D) Representative flow plots (C) and boxplots (D) showing the proportion of classical NK cell subsets defined by expression of CD56 and CD16 in NK cells from healthy, moderate COVID (“Moderate”), and severe COVID (“Severe”) donors. E) Boxplots showing the mean fluorescence intensity (MFI) of activating receptors DNAM-1 and NKG2D, cytotoxic molecules Granzyme B and Perforin, activation markers CD38 and CD69, and proliferation marker Ki-67 in all NK cells from patients across severity groups. F) Boxplots showing the proportion of activated (CD38^+^ CD69^+^) NK cells in each of the three NK cell subsets identified in subfigure 1B across severity groups. G) Boxplots showing expression of CD69 and Ki-67 in the NK cells of fatal COVID-19 cases compared to healthy or hospitalized, non-fatal cases. Significance values for all plots in this figure were determined using an unpaired Wilcoxon rank-sum test with the Bonferroni correction for multiple hypothesis testing.

The NK cells of COVID-19 patients also exhibited changes in their expression of key surface and intracellular molecules (Fig. 2E; Fig S1). Surface expression of the activating receptor NKG2D and the activating co-receptor DNAM-1 were decreased in COVID-19 patients compared to healthy controls, while having greatly increased intracellular expression of the cytotoxic molecules Granzyme B and Perforin and the proliferation marker Ki-67. Markers of activation CD38 and CD69 were more highly expressed on COVID-19 patient NK cells. Notably, Granzyme B, Perforin, and CD38 were all predicted downstream targets of ligands expressed by monocytes in severe COVID-19 (Fig. 1E). These changes were generally conserved in all three NK cell subsets analyzed (CD56^bright^ CD16^lo^, CD56^dim^ CD16^lo^, and CD56^dim^ CD16^hi^) (Fig. S2A-D). However, we found that the proportion of activated NK cells (defined as CD38+ CD69+) increased more substantially in the unconventional CD56^dim^ CD16^lo^ population than in the other subsets (Fig. 1F).

Most analyses of NK cells thus far have not had adequate numbers of fatal COVID-19 samples to assess differences between hospitalized patients who survived their disease course and those that did not. Given that our cohort contained a relatively high number of fatal COVID-19 cases (7), we assessed whether any NK cell markers were differentially expressed between fatal and non-fatal patients. While expression of most markers was not significantly different between fatal and non-fatal cases (Fig. S2E), we identified two markers, CD69 and Ki-67, that were more highly expressed in fatal cases compared to hospitalized but non-fatal cases (Fig. 2G).

The samples from hospitalized COVID-19 patients in this cohort were collected from 2020 to 2021, before the availability of standardized treatment recommendations or outpatient therapeutics such as nirmatrelvir/ritonavir, and the patients were therefore treated with a variety of therapeutics (Fig. 2A). To assess any potential effects of these interventions on NK cell phenotype, we compared NK cells from patients who did not receive each treatment to those who did. In general, we saw no differences in treated versus untreated groups except for those clearly correlated with disease severity (Fig. S3B-G); it is difficult to disentangle effects due to treatment from those due to severity given that interventions are often administered on a severity-dependent basis. However, we did find that the NK cells from the four patients on extracorporeal membrane oxygenation (ECMO) had significantly lower expression of Perforin than those of patients not on ECMO (Fig. S3E).

### COVID-19 patient NK cells exhibit defective killing of SARS-CoV-2-infected and bystander cells

Although the phenotype of severe COVID-19 patient NK cells suggests that they have high cytotoxic potential, other studies have shown that these cells respond poorly to K562 tumor target cells(1, 2, 9, 10). Compared to NK cells from healthy donors, COVID-19 patient NK cells are also worse at reducing viral replication when co-cultured with SARS-CoV-2-infected cells(9, 10). However, no study to date has directly assessed the ability of COVID-19 patient NK cells to kill SARS-CoV-2-infected cells and bystander (uninfected) cells. We previously demonstrated that NK cells are impaired in their ability to lyse SARS-CoV-2-infected target cells due to target cell loss of the ligands for the activating receptor NKG2D(25). We therefore performed killing assays as previously described(25) using NK cells isolated from either healthy donors or COVID-19 patients and A549-ACE2 target cells infected with an mNeonGreen-tagged strain of SARS-CoV-2. By infecting the target cells with an MOI sufficient to result in ∼25% infected cells and utilizing the mNeonGreen fluorescent tag, we were able to differentiate between direct killing of SARS-CoV-2-infected cells and bystander cells. There were no significant differences in the proportion of SARS-CoV-2 NP+ target cells between wells containing NK cells from healthy donors, wells containing NK cells from COVID-19+ donors, and wells containing target cells only (Fig. 3B). We found that COVID-19 patient NK cells were significantly worse at killing bystander A549-ACE2 cells compared to NK cells from healthy donors (Fig. 3C). Additionally, we found that COVID-19 patient NK cells appeared to exhibit a disease severity-dependent defect in their ability to kill SARS-CoV-2-infected cells, although this reduction was not statistically significant (Fig. 3C).

**Fig. 3:**
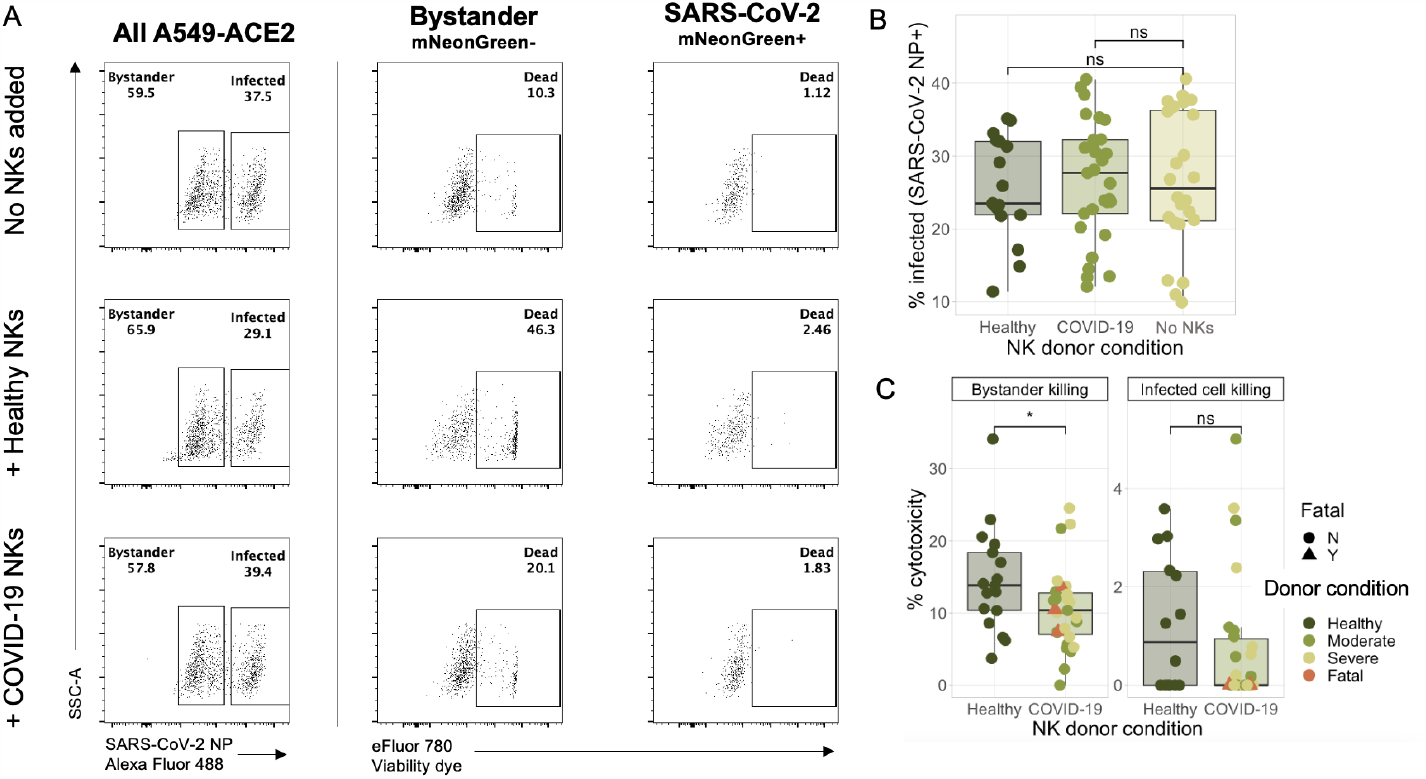
COVID-19 patient NK cells exhibit defective killing of SARS-CoV-2-infected and bystander cells. A) Representative flow plots showing the percentage of infected A549-ACE2 target cells (left) and percentage of dead target cells (center and right) in wells with target cells only (top), healthy NK cells plus target cells (middle), and COVID-19 NK cells plus target cells (bottom). B) Boxplot showing the percentage of infected (SARS-CoV-2 NP+) target cells in wells with healthy or COVID-19+ NK cells versus wells with target cells only. C) Boxplot showing the background-subtracted killing of bystander (left) or SARS-CoV-2-infected (right) target cells after a 3-hour co-culture with NK cells from either healthy or COVID-19+ donors. An effector:target ratio of 10:1 was used. Significance values for all boxplots were determined using a Wilcoxon ranked-sum test.

### Monocytes from severe COVID-19 patients induce activation and proliferation in healthy NK cells

To experimentally validate whether NK-monocyte interactions could indeed underlie NK cell activation in severe COVID-19, we developed an allogeneic co-culture system in which CD14+ monocytes from COVID-19 patients were isolated and co-cultured with NK cells derived from healthy donors. After 2 hours of co-culture, we assessed NK cell phenotype by flow cytometry to determine whether COVID-19 patient monocytes were capable of inducing changes in healthy NK cells (Fig. 4A, Fig. S4-5).

**Figure 4:**
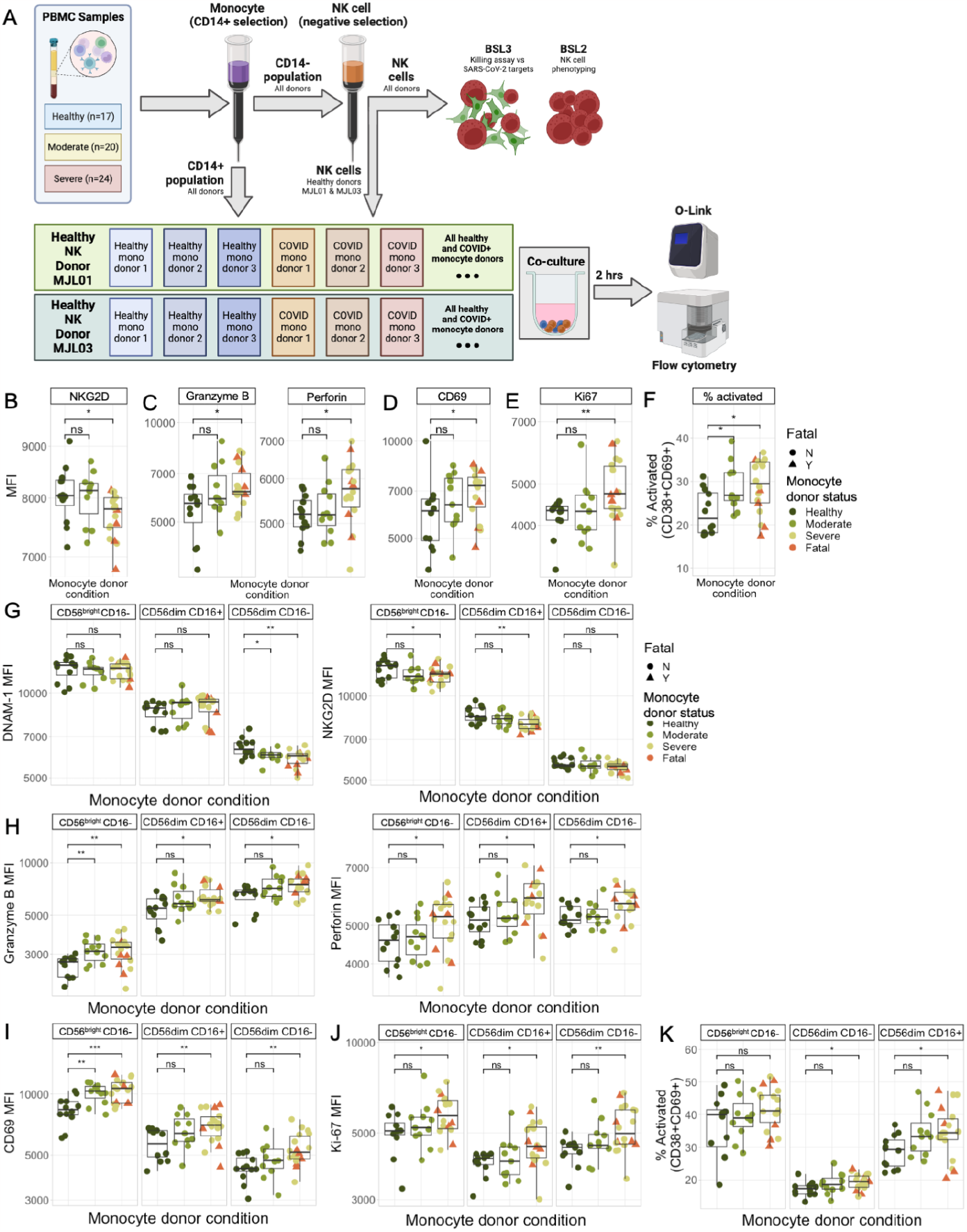
Monocytes from severe COVID-19 patients induce activation and proliferation in healthy NK cells. A) Schematic illustrating the experimental design for allogeneic NK cell-monocyte co-cultures. Briefly, CD14+ monocytes and NK cells were isolated from all donor PBMC. The NK cells from two healthy donors (“MJL01” and “MJL03”) were co-cultured with CD14+ monocytes from allogeneic healthy or COVID-19+ donors for 2 hours in either a round-bottom 96-well plate or a transwell plate. After 2 hours, NK cell phenotypes were assessed via flow cytometry. B-E) Boxplots showing the MFIs of NKG2D (B), Granzyme B and Perforin (C), CD69 (D), and Ki-67 (E) in all NK cells from healthy donors following 2 hour co-culture with allogeneic monocytes. F) Boxplot showing the percent of healthy donor NK cells positive for both CD38 and CD69 following 2 hour co-culture with allogeneic monocytes. G-K) Boxplots showing the MFIs of DNAM-1 and NKG2D (G), Granzyme B and Perforin (H), CD69 (I), and Ki-67 (J) in three distinct subsets of NK cells from healthy donors following 2 hour co-culture with allogeneic monocytes. K) Boxplots showing the percent of healthy donor NK cells from three distinct subsets of NK cells that were positive for both CD38 and CD69 following 2 hour co-culture with allogeneic monocytes. Significance values for all plots in this figure were determined using an unpaired Wilcoxon rank-sum test with the Bonferroni correction for multiple hypothesis testing.

We found that co-culture of healthy NK cells with monocytes from severe COVID-19 patients recapitulated many of the key features of the NK cells from these same severe COVID-19 patients (shown in Fig. 2). Following co-culture with monocytes from severe COVID-19 patients but not moderate patients or healthy donors, healthy NK cells had decreased expression of NKG2D (Fig. 4B). They also had increased expression of the cytotoxic molecules Granzyme B and Perforin (Fig. 4C) as well as the activation and tissue residency marker CD69 (Fig. 4D) and the proliferation marker Ki-67 (Fig. 4E). Finally, NK cells co-cultured with monocytes from both moderate and severe COVID-19 patients had a higher proportion of activated (CD38+ CD69+) cells (Fig. 4F).

As observed in the NK cells of COVID-19 patients, most of the changes induced in healthy NK cells by COVID-19 patient monocytes were conserved across the three NK cell subsets examined (Fig. 4G-K). Expression of Granzyme B, Perforin, Ki-67, and CD69 was increased in CD56^bright^ CD16^lo^, CD56^dim^ CD16^lo^, and CD56^dim^ CD16^hi^ NK cells (Fig. 4H-K). However, expression of DNAM-1 and NKG2D was affected differentially in different subsets: DNAM-1 expression was only significantly decreased in unconventional CD56^dim^ CD16^lo^ cells, while NKG2D was only downregulated in the other two subsets (CD56^bright^ CD16^lo^ and CD56^dim^ CD16^hi^) (Fig. 4G). Together, these data indicate that monocytes from COVID-19 patients are sufficient to induce activation and proliferation of NK cells as well as downregulation of NKG2D and DNAM-1.

### Monocyte-induced CD56^dim^ NK cell activation in severe COVID-19 occurs primarily through contact-dependent mechanisms

To determine mechanisms by which monocytes activate NK cells, we examined whether activation occurred through contact-dependent or contact-independent mechanisms. We performed our co-culture experiments using either a direct culture system (as shown in Fig. 4) or a transwell culture system with monocytes on the bottom and NK cells in a transwell insert to abrogate direct contact. We first analyzed the total population of NK cells following either direct or transwell co-culture with monocytes (Fig. 5A, Fig. S6A) and both CD16+ and CD16-CD56^dim^ NK cells, which typically make up >95% of peripheral blood NK cells (Fig. 5B-C, Fig. S6C-D). While NK cells that underwent direct co-culture with monocytes from severe COVID-19 patients exhibited increases in cytotoxic molecule expression, proliferation, and activation, we found that all of these statistically significant increases in marker expression were abrogated in the NK cells that instead underwent transwell co-culture (Fig. 5A, Fig. S6). These results suggest that the activated, proliferative NK cell phenotype induced by co-culture with monocytes from severe COVID-19 patients is primarily mediated by direct cell-cell contacts in CD56^dim^ NK cells, although small trending increases in Granzyme B, Ki-67, and co-expression of CD38 and CD69 imply that there may be role for soluble factors in this activation as well.

**Figure 5:**
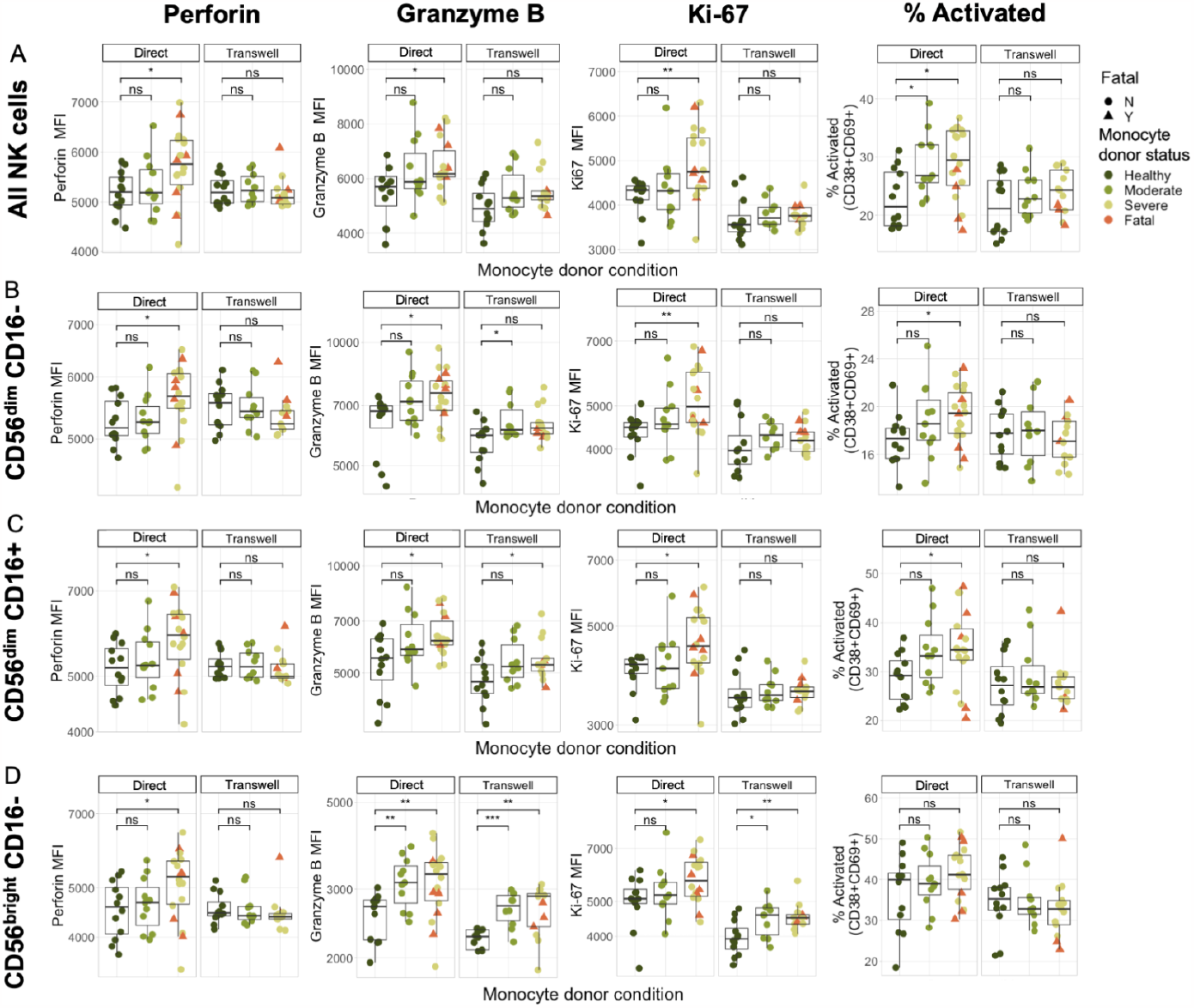
CD56^bright^ NK cells are activated by monocytes through contact-independent mechanisms while CD56^dim^ NK cells are activated primarily through contact-dependent mechanisms. A-D) Boxplots showing the MFI of Perforin, Granzyme B, and Ki-67 and the percentage of NK cells positive for CD38 and CD69 across severity conditions in normal (round-bottom 96 well plate) cultures and transwell cultures. Each row shows the expression of these markers in a different population of NK cells: total NK cells (A); CD56^dim^ CD16-NK cells (B); CD56^dim^ CD16+ NK cells (C); or CD56^bright^ CD16-NK cells (D).

### CD56^bright^ NK cells are activated through soluble interactions with monocytes from severe COVID-19 patients

Although CD56^dim^ NK cells were only minimally activated in a transwell setting, we found that CD56^bright^ CD16-NK cells had strongly increased expression of Granzyme B and Ki-67 after transwell co-culture with monocytes from severe COVID-19 patients (Fig. 5D), although these cells did not undergo an increase in Perforin expression in a transwell setting (Fig. 5D). These results suggest that the mechanisms by which monocytes from severe COVID-19 patients activate NK cells differ between subsets of NK cells, with CD56^bright^ CD16^-^ cells being more strongly affected by soluble factors and CD56^dim^ NK cells being more strongly affected by direct cell-cell interactions.

### Pro-inflammatory and pro-chemotactic cytokines contribute to the NK cell phenotype

We next sought to identify soluble factors that may have induced activation of CD56^bright^ CD16^-^ NK cells in the presence of monocytes from severe COVID-19 patients. We performed O-link analysis using the Inflammation Panel to assay 92 soluble markers including cytokines, chemokines, and growth factors on the supernatants from our transwell co-culture experiments. Due to limited cell numbers, we were not able to evaluate monocytes in the absence of NK cells. Of the 92 analytes measured, 58 were detected in at least one sample measured (Fig. 6A). We performed multidimensional scaling analysis using the 58 markers detected, revealing almost complete separation between severe COVID-19 patient samples and healthy donors, with moderate COVID-19 patient samples occupying an intermediate space (Fig. 6B). Interestingly, the three fatal donor samples included in this analysis clustered more closely with the healthy donor samples than with the severe, non-fatal donor samples (Fig. 6B). The separation between severe and healthy donors in multidimensional space was driven by increased expression of a wide variety of analytes in the severe patient samples, including MMP-1, MCP-3, CXCL5, IL-18, CXCL1, VEG-F-A, and Latency-associated peptide TGFβ1 (LAP TGF B1) (Fig. 6C).

**Figure 6:**
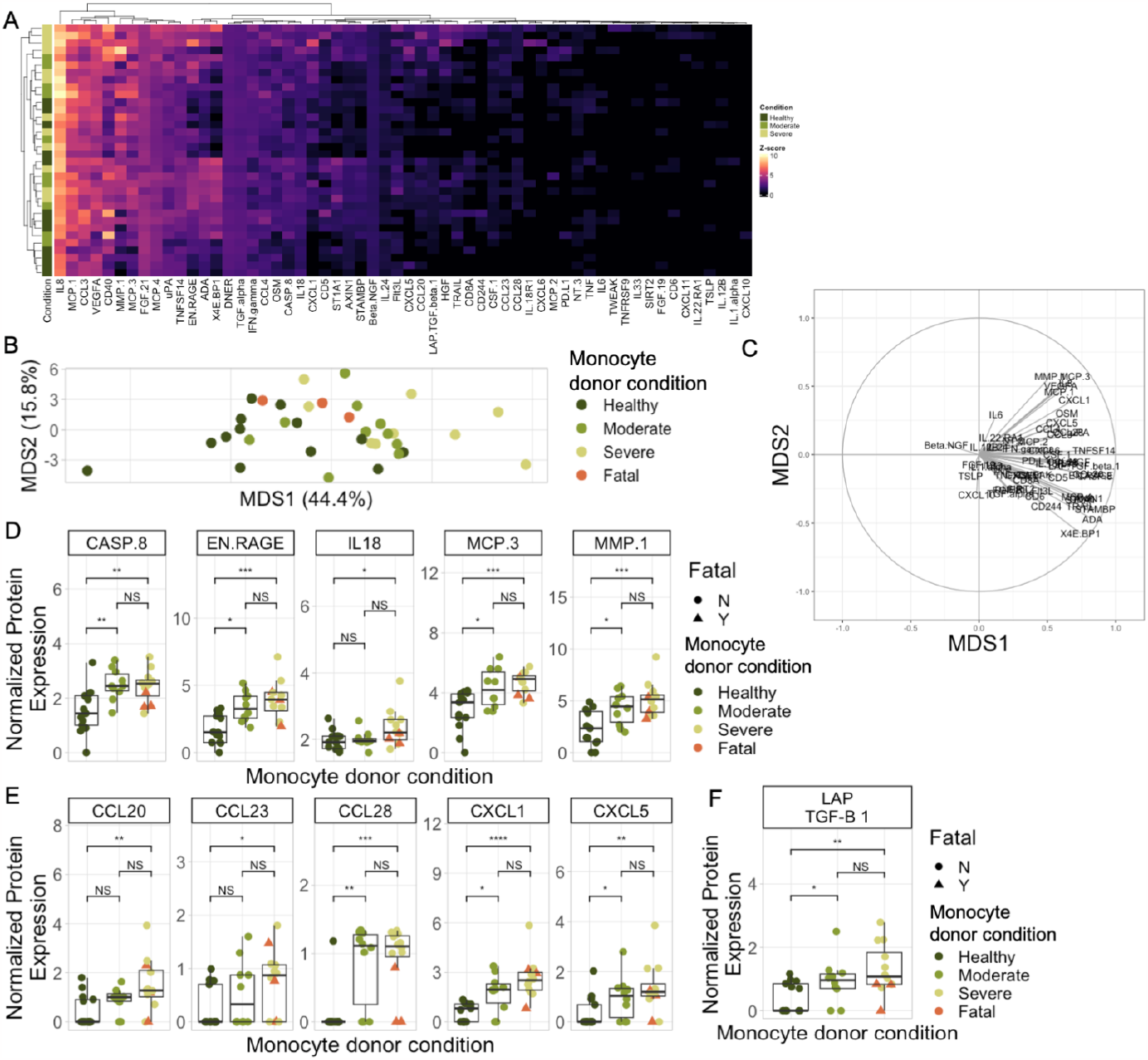
Monocytes from COVID-19 patients induce an inflammatory and pro-chemotactic cytokine environment. A) Heatmap showing normalized protein expression of the 58 analytes from the O-link Inflammation Panel that were detected in at least one sample. B) Multidimensional scaling (MDS) plot. C) Circle plot. D-F) Boxplots showing the normalized protein expression of various analytes in the O-link Inflammation Panel across monocyte donor severity conditions.

Interrogating expression of individual markers, we found five notable pro-inflammatory analytes (Caspase 8, EN-RAGE, IL-18, MCP-3, and MMP-1) that were present at higher concentrations in the supernatants of severe COVID-19 patient monocyte cultures compared to healthy monocyte cultures (Fig. 6D, Fig. S8). Cultures containing monocytes from severe COVID-19 patients also had significantly elevated levels of many chemokines that are known to regulate the immune response to infectious disease (Fig. 6E). In addition to the observed increase in expression of inflammatory analytes and chemokines, we also found a severity-dependent increase in LAP TGFβ1 (Fig. 6F), which is used as a biomarker for active TGFβ1(26, 27). TGFβ1 itself could not be accurately measured in these samples because active TGFβ1 detection assays cross-react with the fetal bovine serum (FBS) present in the cell culture medium(27). As previously noted, high levels of TGFβ1 in the serum of severe COVID-19 patients have been shown to induce NK cell dysfunction in COVID-19(9).

## DISCUSSION

Although it has been well-established that peripheral NK cells from severe COVID-19 patients take on a unique phenotype and are highly dysfunctional, the mechanisms underlying these changes were not previously well-understood. Other studies have suggested that this phenotype may partially result from type I interferon signaling(4, 9, 10) and TGFβ in the serum of COVID-19 patients has been shown to have an inhibitory effect on NK cells(9). Crosstalk between NK cells and other immune cells is a key regulator of the NK cell response in other infections(14, 15, 19, 28, 29), leading us to interrogate how NK cell interactions with other cell types underlie NK cell activation and dysfunction in COVID-19. Here, we performed computational analyses showing that NK cells interact strongly with monocytes in severe COVID-19. We then validated experimentally that monocytes from severe COVID-19 patients can induce an activated phenotype in healthy NK cells, similar to the phenotype observed in primary NK cells from severe COVID-19 patients. We further demonstrated that these interactions occur via both contact-dependent and contact-independent mechanisms. Our results collectively illustrate the importance of NK cell communication with other peripheral immune cells in severe COVID-19 and explore a novel mechanism of NK cell activation in this setting.

All three major subsets of NK cells analyzed (CD56^bright^ CD16^-^, CD56^dim^ CD16^-^, and CD56^dim^ CD16^+^) underwent significant phenotypic changes following direct co-culture with monocytes from severe COVID-19 patients. However, the mechanisms underlying the changes in each of these subsets appears to differ; nearly all of the changes in CD56^dim^ NK cells following direct co-culture were abrogated when the co-culture was performed in a transwell setting, while CD56^bright^ NK cells were activated in both transwell and direct settings. These results imply that CD56^dim^ NK cells are activated primarily through contact-dependent mechanisms. Indeed, results from our previous work suggested that NK cells may receive activation through the expression of ligands for NKG2D and DNAM-1 on monocytes in severe COVID-19 patients(3). This hypothesis is further supported by the fact that downregulation of DNAM-1 on CD56^dim^ CD16^-^ NK cells and NKG2D on CD56^dim^ CD16^+^ NK cells was only observed in direct and not transwell cultures in this study (Fig. S7). Both NKG2D and DNAM-1 can be internalized following ligation, leading to loss of expression on the cell surface(30, 31). Our use of the recently-developed computational technique Scriabin to infer cell-cell communication between NK cells and other peripheral immune cells at the single-cell level also provided candidate contact-dependent interactions that may underlie activation and dysfunction of CD56^dim^ NK cells by monocytes in severe COVID-19. Notably, many of the interactions predicted by Scriabin to be upregulated in severe COVID-19 are interactions between class I MHC molecules and KIR or LILRB family receptors, which typically have an inhibitory effect on NK cell cytotoxicity(32, 33). The activating and inhibitory signals received by CD56^dim^ NK cells from monocytes in severe COVID-19 may therefore explain these cells’ activated yet dysfunctional state.

While CD56^dim^ NK cell activation by monocytes was largely abrogated in transwell co-cultures, Granzyme B and Ki-67 in CD56^bright^ NK cells were strongly upregulated in transwell co-cultures (Fig. 5B). A plausible explanation for this observation is the fact that CD56^bright^ NK cells typically express much higher levels of cytokine and chemokine receptors than their CD56^dim^ counterparts and may therefore experience stronger activation by the suite of proinflammatory mediators present in the supernatants of cultures containing severe COVID-19 patient monocytes. Three of these analytes (Caspase-8, EN-RAGE, and IL-18) are all associated with the NLRP3 inflammasome(34–37). In addition to its role in inducing cell death, Caspase-8 also has a non-cytotoxic role in modulating NK and CD8^+^ T cell responses to viral infection(38). RAGE, the receptor for EN-RAGE (S100A12), is expressed on human(39) and murine(40) NK cells. S100A8/9 have been shown to activate NK cells through RAGE(40), although the ability of S100A12 to activate NK cells through RAGE has not yet been directly assessed. Monocyte chemoattractant protein 3 (MCP-3), also known as CCL7, induces not only lymphocyte chemotaxis but also NK cell activation(41). CD56^bright^ NK cells express increased levels of the receptors for MCP-1(42, 43) and IL-18R(44, 45) as well as TGFβ receptor 2 and TGFβ receptor 3(46). Conversely, CD56^dim^ NK cells express higher levels of the killer immunoglobulin-like receptors (KIRs), whose contact-dependent interactions with MHC class I on monocytes were identified by Scriabin as being significantly upregulated in severe COVID-19 (Fig. 1D).

The results of this study generate new insights into NK cell and monocyte trafficking in severe COVID-19. We found that co-cultures of severe COVID-19 patient monocytes and healthy NK cells had significantly increased concentrations of multiple chemokines compared to co-cultures of healthy donor monocytes with healthy NK cells. The upregulated chemokines include CCL23(47), CCL28(48), CXCL1(49), and CXCL5(50), all of which are involved in monocyte chemotaxis. CCL23 also stimulates secretion of matrix metalloproteinases in monocytes(47). CCL20 is not chemoattractive to monocytes, but does induce chemotaxis in NK cells and other lymphocytes(51, 52). Notably, the cells used in these experiments were derived from blood rather than from the site of infection in the airways; therefore, the role of these chemokines in recruiting immune cells to participate in the immune response to SARS-CoV-2 in this setting is unclear. However, in addition to examining chemokine secretion, we also assessed expression of CD69 on NK cells, which is a marker of tissue residency(53) and actively aids NK cell retention in tissue(54). CD69 is upregulated on the NK cells of severe COVID-19 patients(3, 5, 9, 55, 56) and was significantly upregulated following co-culture of healthy NK cells with severe COVID-19 patient monocytes in our study (Fig. 4B, Fig. S8). NK cells have also been shown to traffick to the site of infection in severe COVID-19(6, 11–13, 42). Therefore, interactions between NK cells and monocytes in the periphery may drive CD69 expression that leads to NK cell retention in the lungs during severe COVID-19.

Activation of NK cells by monocytes has been documented in several other infectious diseases, but overall, relatively little is known about monocyte-NK cell crosstalk in viral infection in comparison to NK cell communication with cell subsets such as dendritic cells and T cells(28). Most work thus far has focused on the ability of NK cells to lyse virally-infected monocytes and macrophages(18, 57, 58). The small body of work examining crosstalk between NK cells and uninfected monocytes has centered around monocyte secretion of cytokines that are known to modulate NK cell responses, including IL-12(59) and IL-18(57). Our work here describes both soluble and contact-dependent interactions that take place between NK cells and uninfected monocytes away from the site of infection and influence NK cell phenotype. These insights further our understanding of immune polyphony in the setting of infectious disease, which in turn informs the designs of critical vaccines and therapeutics seeking to modulate the immune response.

## METHODS

### Cohort

Samples from hospitalized COVID-19 patients were obtained from an observational cohort study of hospitalized COVID-19 patients at UCLA. All participants signed informed consent to participate and the study was approved by the UCLA Institutional Review Board (#20-000473). Patients were recruited from two UCLA Health hospitals in Los Angeles, CA. Inclusion criteria included hospitalization for COVID-19, age greater than 18, and confirmed positive SARS-CoV-2 RT-PCR within 72 hours of admission. Exclusion criteria included pregnancy, hemoglobin less than 8 g/dL, or inability to provide informed consent. Upon enrollment blood samples, nasopharyngeal swab, and saliva were collected throughout hospitalization up to 6 weeks. Demographic and clinical data, including therapeutics, were collected from the electronic medical record. Clinical severity was scored using the NIAID 8-point ordinal scale(60): 1, not hospitalized and no limitations; 2, not hospitalized but with limitations; 3, hospitalized no supplemental oxygen or ongoing medical care; 4, hospitalized no supplemental oxygen but with ongoing medical care; 5, hospitalized with supplemental oxygen; 6, hospitalized with non-invasive ventilation or high-flow oxygen; 7, hospitalized with invasive mechanical ventilation or ECMO; 8, death. For this study, mild COVID-19 included ordinal scale 3-4, moderate COVID-19 included ordinal scale 5, and severe COVID-19 included ordinal scale 6-7. Samples included in this study were collected from April 2020 through February 2021.

### Monocyte Isolation

Cryopreserved PBMC from COVID-19-positive and healthy donors were thawed at 37 C and washed with RPMI supplemented with 10% FBS (RP10) to remove freezing medium. Cells from each donor were magnetically fractionated into CD14+ and CD14-populations using the Miltenyi MACS Human CD14+ MicroBead isolation kit (Miltenyi, cat. no. 130-050-201) according to the manufacturer’s instructions. CD14+ cells were then set aside in an incubator (37 C, 5% CO_2_) until the start of co-culture while CD14-cells were used for subsequent NK cell isolation.

### NK cell isolation and activation

NK cells were isolated from CD14-cells from healthy donors and COVID-19 patients using the Miltenyi MACS Human NK Cell Isolation Kit (Miltenyi, cat. no. 130-092-657) according to the manufacturer’s instructions. 10% of NK cells from healthy and COVID-19-positive donors were set aside for phenotyping by flow cytometry. The remaining NK cells were transferred to a round-bottom 96-well plate and resuspended in complete RPMI supplemented with 25 ng/mL (250 IU/mL) rhIL-2 (R&D Systems, cat. no. 202-IL-010), then placed in a 37C CO_2_ incubator for 12-16 hours. After incubation, the cells were washed twice to remove IL-2, counted, and resuspended in fresh RP10 before being transferred to BSL3 facilities for killing assays.

### NK cell phenotyping

As described above, a sample of NK cells from healthy donors and COVID-19 patients were taken for phenotyping by flow cytometry. NK cells were washed in PBS and stained with eFluor 780 Fixable Viability Dye (eBioscience, cat. no. 65-0865-14) for 20 minutes. Cells were then washed in FACS buffer (PBS supplemented with 2% FBS) and stained for 30 minutes at room temperature with a panel of antibodies against surface antigens. Stained NK cells were washed, fixed for 15 minutes in 4% paraformaldehyde (PFA) (EIS, cat. no. 15710), and permeabilized (BD Biosciences, Cat. 340973). Permeabilized cells were stained with a panel of antibodies against intracellular proteins, then washed and analyzed on a Cytek Aurora spectral cytometer.

### Cell lines

A549-ACE2 were a gift from Ralf Bartenschlager and were confirmed to be mycoplasma-free. A549-ACE2 cells were maintained in DMEM supplemented with 10% FBS and passaged every 2-3 days. Cell cultures were discarded and new cells were thawed after 25 passages.

### Infection of A549-ACE2 with SARS-CoV-2

The day prior to infection, A549-ACE2 cells were seeded at a density of 100K cells/well in a 12-well plate. On the day of infection, cells were brought into the BSL3 and washed once with PBS to remove excess serum. PBS was then removed and mNeon Green SARS-CoV-2 was added in DMEM supplemented with 2% FBS (“D2”) at an MOI of 0.5 (final volume of 150 uL per well). The plate was rocked for 1 hour at 37C, after which time the virus was washed off with PBS and 0.5 mL D2 was added to each well. Plates were placed back in an incubator for 48 hours before being harvested for use in killing assays.

### Flow cytometry-based killing assay

The morning of the killing assay, IL-2-activated NK cells were counted and brought into the BSL3. SARS-CoV-2-infected A549-ACE2s were washed with PBS, harvested using TrypLE, and counted before being resuspended in fresh RP10. Target cells and NK cells were plated in V-bottom 96-well plates at an effector:target (E:T) ratio of 10:1. The plate containing target cells and NK cells was spun down for 1 minute at 1000 RPM to bring cells together, then placed in the 37 C incubator for 3 hours. After 3 hours, cells were washed with PBS and stained with eFluor 780 Fixable Viability Dye for 25 minutes. Cells were then washed in PBS and fixed for 30 minutes in 4% PFA before being transferred to fresh tubes, decontaminated, and removed from the BSL3 and analyzed on a Cytek Aurora spectral cytometer.

### Allogeneic NK cell/monocyte co-culture

Purified CD14+ cells in RP10 were added to a round-bottom 96 well plate or the bottom of a 24-well transwell plate for a final concentration of 1.5x10^6^ CD14+ cells/mL. Two replicates were plated for each monocyte donor; each of these replicates then received NK cells from one of two healthy donors (MJL01 or MJL03) at a final concentration of 0.75x10^6^ NK cells/mL. NK cells were added directly to monocytes in round-bottom 96 well plates or to the top of a 0.4 um transwell insert in transwell plates. Both culture systems had a monocyte:NK ratio of 2:1 and the final concentration of cells in media was kept consistent between the two culture systems. Once the cells had been added, 96 well culture plates were spun down for 1 minute at 1000 RPM to bring the cells together. Spun-down 96 well plates and transwell culture plates were then placed in a 37 C incubator for 2 hours. After 2 hours, NK cells from the transwell inserts were collected and transferred to a fresh 96-well plate. NK cells from all cultures were then stained for flow cytometry and analyzed in the manner described above (under “NK cell phenotyping”). The transwell and direct cultures for each donor were performed simultaneously to minimize batch effects.

### O-link

After the NK cells were harvested from the transwell cultures, the remaining transwell culture supernatants were saved for O-link analysis. The supernatants were collected in microcentrifuge tubes and centrifuged to remove any cells and cell debris in the sample. Once clarified, the supernatants were transferred to fresh tubes and frozen at -80C until analyzed. Samples did not undergo any freeze-thaw cycles other than when they were thawed for final analysis.

### Scriabin analysis

scRNA-seq data from Wilk et al. 2021(3) was first passed through a denoising algorithm (“ALRA”), which uses low-rank matrix approximation to impute expression levels of lowly-expressed genes, thereby partially alleviating the sparsity of the gene expression matrix(61). Scriabin was then used to identify sets of highly co-expressed ligand-receptor pairs and group them into Interaction Programs (“IPs”) whose expression could be compared between sample conditions and cell types.

Scriabin was also used to generate a cell-cell interaction matrix (“CCIM”) from ALRA-denoised data, which consists of expression levels of all ligand-receptor pairs for each pair of cells within the dataset. In creating this matrix, Scriabin utilizes NicheNet to weight expression of each LR pair by expression of genes downstream of said pair in order to identify biologically-active LR pairs. Creation of a CCIM for all monocytes and NK cells in our dataset allowed us to interrogate weighted expression of individual LR edges across disease severity conditions. To discover specific LR pairs that may define monocyte-NK cell interactions in severe COVID-19, we applied the Seurat function FindMarkers to the CCIM to identify other differentially-expressed edges in these patients compared to healthy controls.

### Quantification and statistical analysis

Flow cytometry data visualization was performed using FlowJo v10.7.1. Figures were generated in R using the *ggplot2, Seurat*, and *Scriabin* packages. Colors for figures were generated using the *NatParksPalettes* package. Statistical analyses were performed as described in figure legends and plotted using the R *ggpubr* package.

## ACKNOWLEDGEMENTS

We are very grateful to the donors who provided peripheral blood for these experiments and to their families. We thank Dr. Pei-Yong Shi for the kind gift of icSARS-CoV-2/WA-01-mNeonGreen. We thank Dr. Jaishree Garhyan for her assistance in Stanford’s BSL3 facilities. We are grateful to Minne Lee and Matthew Kaufmann for their insightful commentary on the work. Figure illustrations were created using BioRender.com. Data analysis and visualization was performed using R.

Funding was provided by the Bill & Melinda Gates Foundation OPP1113682 (to C.A.B.), Chan Zuckerberg Biohub (to C.A.B.), the Burroughs Wellcome Fund Project 1016687 (to C.A.B.), a Stanford Chem-H/Innovative Medicine Accelerator COVID-19 Response Award (to C.A.B.), and the National Institutes of Health (U19 AI057229). Fellowship and training support was from National Institutes of Health (T32 AI00729037 and F31 AI172311-01 to M.J.L.) and the Stanford Pandemic Preparedness Hub (to T.R.B). C.A.B. is an investigator of the Chan Zuckerberg Biohub.

