## Supplementary Table 1 for "NK cell-monocyte crosstalk underlies NK cell activation in severe COVID-19"

| lr_pair | name | severe_pval | healthy_pval | severe_conne | healthy_connectivity |
|---|---|---|---|---|---|
| DLL1=NOTCH: | IP-1 | 0.0001 | 0.1327 | 0.59040003 | 0.75106466 |
| EFNA1=EPHB1 | IP-1 | 0.0001 | 0.1327 | 0.57398927 | 0.76229272 |
| ERBB2=ERBB3 | IP-1 | 0.0001 | 0.1327 | 0.47811601 | 0.68964519 |
| IGF2=IGF1R | IP-1 | 0.0001 | 0.1327 | 0.59468691 | 0.78257292 |
| IGF2=IGF2R | IP-1 | 0.0001 | 0.1327 | 0.57535173 | 0.76377911 |
| IL12A=IL12RB | IP-1 | 0.0001 | 0.1327 | 0.58791747 | 0.76997123 |
| IL17A=IL17RA | IP-1 | 0.0001 | 0.1327 | 0.56761063 | 0.76754829 |
| IL17A=IL17RC | IP-1 | 0.0001 | 0.1327 | 0.56737004 | 0.76587497 |
| IL25=IL17RB | IP-1 | 0.0001 | 0.1327 | 0.55656491 | 0.76930055 |
| NCAM1=FGFR | IP-1 | 0.0001 | 0.1327 | 0.53454417 | 0.70342913 |
| RLN2=RXFP2 | IP-1 | 0.0001 | 0.1327 | 0.5845312 | 0.78204393 |
| SEMA7A=PLXI | IP-1 | 0.0001 | 0.1327 | 0.65993156 | 0.82980314 |
| XCL1=XCR1 | IP-1 | 0.0001 | 0.1327 | 0.56205717 | 0.75659613 |
| CALCB=RAMP | IP-10 | 0.0182 | 0.1449 | 0.54482358 | 0.75821038 |
| COL18A1=GP( | IP-10 | 0.0182 | 0.1449 | 0.56885274 | 0.76439907 |
| FGF5=FGFR1 | IP-10 | 0.0182 | 0.1449 | 0.59523864 | 0.75534926 |
| GDF6=BMPR1 | IP-10 | 0.0182 | 0.1449 | 0.55417639 | 0.7581147 |
| GDF6=BMPR2 | IP-10 | 0.0182 | 0.1449 | 0.59366252 | 0.76771986 |
| ADAM17=NO | IP-13 | 0 | 0 | 0.93244589 | 0.91031936 |
| ADAM9=ITGA | IP-13 | 0 | 0 | 0.83387244 | 0.95552805 |
| ALCAM=CD6 | IP-13 | 0 | 0 | 0.59957352 | 0.84002104 |
| BAG6=NCR3 | IP-13 | 0 | 0 | 0.52679041 | 0.65955997 |
| CXCL16=CXCR | IP-13 | 0 | 0 | 0.62385603 | 0.85146955 |
| ICAM1=ITGAL | IP-13 | 0 | 0 | 0.99784278 | 0.92810535 |
| ICAM1=ITGAN | IP-13 | 0 | 0 | 0.977666 | 0.95039997 |
| ICAM1=ITGB2 | IP-13 | 0 | 0 | 1 | 0.98149421 |
| IL15=IL2RB | IP-13 | 0 | 0 | 0.56477991 | 0.76431507 |
| IL15=IL2RG | IP-13 | 0 | 0 | 0.7447103 | 0.83650527 |
| INSR=ADRB2 | IP-13 | 0 | 0 | 0.93664696 | 0.94015887 |
| JAG1=NOTCH: | IP-13 | 0 | 0 | 0.87470324 | 0.92436765 |
| LGALS9=HAV( | IP-13 | 0 | 0 | 0.93860412 | 0.87878614 |
| NRG1=ERBB2 | IP-13 | 0 | 0 | 0.5466478 | 0.75337675 |
| PVR=CD226 | IP-13 | 0 | 0 | 0.60952202 | 0.8080848 |
| SIRPA=CD47 | IP-13 | 0 | 0 | 0.75251255 | 0.90748531 |
| TGFB1=TGFBF | IP-13 | 0 | 0 | 0.54219406 | 0.68103431 |
| TIMP1=CD63 | IP-13 | 0 | 0 | 0.83792773 | 0.94487467 |
| ANGPT1=TIE1 | IP-17 | 0.0002 | 0.0005 | 0.53704093 | 0.77046068 |
| APOE=LRP8 | IP-17 | 0.0002 | 0.0005 | 0.70059912 | 0.79911382 |
| APOE=VLDLR | IP-17 | 0.0002 | 0.0005 | 0.61285988 | 0.77908389 |
| BMP4=BMPR: | IP-17 | 0.0002 | 0.0005 | 0.61344759 | 0.78911158 |
| BMP4=BMPR: | IP-17 | 0.0002 | 0.0005 | 0.646575 | 0.80084393 |
| BMP4=BMPR: | IP-17 | 0.0002 | 0.0005 | 0.69038242 | 0.81661979 |
| GDF9=BMPR1 | IP-17 | 0.0002 | 0.0005 | 0.57152717 | 0.78369717 |
| IL23A=IL12RB | IP-17 | 0.0002 | 0.0005 | 0.54208157 | 0.78560253 |
| NCR3LG1=NCI | IP-17 | 0.0002 | 0.0005 | 0.55378099 | 0.69972482 |
| POMC=MC1R | IP-17 | 0.0002 | 0.0005 | 0.62361958 | 0.74033156 |
| CCL2=CCR5 | IP-18 | 0.7837 | 0.0328 | 0.7719236 | 0.84308466 |
| CCL3=CCR5 | IP-18 | 0.7837 | 0.0328 | 0.80923394 | 0.87668448 |
| CCL3L3=CCR5 | IP-18 | 0.7837 | 0.0328 | 0.81800447 | 0.87444089 |

| | | | | | |
|---|---|---|---|---|---|
| IFNL2=IL10RB | IP-18 | 0.7837 | 0.0328 | 0.53114612 | 0.74971285 |
| NPW=NPBWR | IP-18 | 0.7837 | 0.0328 | 0.46884421 | 0.00724511 |
| PLG=F2RL1 | IP-18 | 0.7837 | 0.0328 | 0.71049921 | 0.81021998 |
| TIMP2=ITGA3 | IP-18 | 0.7837 | 0.0328 | 0.57981617 | 0.79290749 |
| CCL8=CCR1 | IP-20 | 0.9672 | 0.0288 | 0.71362018 | 0.80929468 |
| CCL8=CCR2 | IP-20 | 0.9672 | 0.0288 | 0.70129007 | 0.8041171 |
| GAL=GALR2 | IP-20 | 0.9672 | 0.0288 | 0.49369956 | 0.21053728 |
| INHBB=ACVR1 | IP-20 | 0.9672 | 0.0288 | 0.71683626 | 0.74999524 |
| INHBB=ACVR2 | IP-20 | 0.9672 | 0.0288 | 0.69419724 | 0.74336803 |
| ADAM17=ERB | IP-22 | 0.0018 | 0.0073 | 0.63548858 | 0.80665809 |
| DLL3=NOTCH | IP-22 | 0.0018 | 0.0073 | 0.59049032 | 0.77253516 |
| DLL3=NOTCH | IP-22 | 0.0018 | 0.0073 | 0.58866243 | 0.78086352 |
| DLL3=NOTCH | IP-22 | 0.0018 | 0.0073 | 0.57186626 | 0.76469912 |
| DLL3=NOTCH | IP-22 | 0.0018 | 0.0073 | 0.58613242 | 0.77761994 |
| ERBB2=ERBB4 | IP-22 | 0.0018 | 0.0073 | 0.5132975 | 0.74050055 |
| EREG=ERBB4 | IP-22 | 0.0018 | 0.0073 | 0.63228809 | 0.80173436 |
| HBEGF=ERBB | IP-22 | 0.0018 | 0.0073 | 0.64832957 | 0.8077896 |
| IFNW1=IFNAR | IP-22 | 0.0018 | 0.0073 | 0.52000497 | 0.71515652 |
| IL13=IL13RA1 | IP-22 | 0.0018 | 0.0073 | 0.57766948 | 0.75151627 |
| IL13=IL4R | IP-22 | 0.0018 | 0.0073 | 0.55769607 | 0.74701492 |
| IL17F=IL17RA | IP-22 | 0.0018 | 0.0073 | 0.52586859 | 0.46537421 |
| NRG1=ERBB4 | IP-22 | 0.0018 | 0.0073 | 0.61740903 | 0.80568999 |
| NRG2=ERBB3 | IP-22 | 0.0018 | 0.0073 | 0.56548219 | 0.7577049 |
| NRG4=ERBB4 | IP-22 | 0.0018 | 0.0073 | 0.59713592 | 0.77668256 |
| BMP2=ACVR2 | IP-23 | 0.0001 | 0 | 0.60719046 | 0.79651723 |
| BMP2=ACVR2 | IP-23 | 0.0001 | 0 | 0.59338912 | 0.79520236 |
| BMP2=BMPR | IP-23 | 0.0001 | 0 | 0.57026262 | 0.7811106 |
| BMP2=BMPR | IP-23 | 0.0001 | 0 | 0.58563796 | 0.78767635 |
| BMP2=BMPR | IP-23 | 0.0001 | 0 | 0.60117159 | 0.7995261 |
| COL1A1=DDR | IP-23 | 0.0001 | 0 | 0.54438167 | 0.7610877 |
| EGFR=ERBB2 | IP-23 | 0.0001 | 0 | 0.51841382 | 0.74827851 |
| IL12A=IL12RB | IP-23 | 0.0001 | 0 | 0.58791747 | 0.76997123 |
| PDGFD=PDGF | IP-23 | 0.0001 | 0 | 0.50020011 | 0.63150855 |
| ADCYAP1=VIP | IP-24 | 0 | 0 | 0.59222203 | 0.74525893 |
| ADM=CALCRL | IP-24 | 0 | 0 | 0.81743982 | 0.90926088 |
| ANXA1=FPR1 | IP-24 | 0 | 0 | 0.72963551 | 0.90819847 |
| APOB=LDLR | IP-24 | 0 | 0 | 0.58162229 | 0.76671305 |
| APP=CD74 | IP-24 | 0 | 0 | 0.82911652 | 0.97071632 |
| APP=LRP1 | IP-24 | 0 | 0 | 0.80511031 | 0.96626967 |
| CCL14=CCR1 | IP-24 | 0 | 0 | 0.55973366 | 0.75482269 |
| CCL14=CCR5 | IP-24 | 0 | 0 | 0.57607103 | 0.7512097 |
| CCL2=CCR1 | IP-24 | 0 | 0 | 0.8257071 | 0.88863855 |
| CCL2=CCR2 | IP-24 | 0 | 0 | 0.80012491 | 0.87813081 |
| CCL3=CCR1 | IP-24 | 0 | 0 | 0.88513085 | 0.94388376 |
| CCL3L3=CCR1 | IP-24 | 0 | 0 | 0.91084822 | 0.94075911 |
| CCL5=CCR1 | IP-24 | 0 | 0 | 0.46258815 | 0.75415386 |
| CSF1=CSF1R | IP-24 | 0 | 0 | 0.75004398 | 0.81006548 |
| CTF1=IL6ST | IP-24 | 0 | 0 | 0.62836656 | 0.77563209 |
| EDN1=EDNRB | IP-24 | 0 | 0 | 0.75299802 | 0.77161542 |
| EFNB1=EPHB2 | IP-24 | 0 | 0 | 0.86682713 | 0.88139107 |

| | | | | | |
|---|---|---|---|---|---|
| ERBB3=NRG1 | IP-24 | 0 | 0 | 0.57104526 | 0.78984064 |
| F12=CD93 | IP-24 | 0 | 0 | 0.73361191 | 0.8310069 |
| GAS6=MERTK | IP-24 | 0 | 0 | 0.83377044 | 0.85036214 |
| HBEGF=CD9 | IP-24 | 0 | 0 | 0.74233772 | 0.88890171 |
| ICAM1=ITGAM | IP-24 | 0 | 0 | 0.977666 | 0.95039997 |
| ICAM1=ITGB2 | IP-24 | 0 | 0 | 1 | 0.98149421 |
| IGF2=INSR | IP-24 | 0 | 0 | 0.60109031 | 0.79364999 |
| IL10=IL10RB | IP-24 | 0 | 0 | 0.87909324 | 0.88338511 |
| IL15=IL15RA | IP-24 | 0 | 0 | 0.94535368 | 1 |
| IL1A=IL1RAP | IP-24 | 0 | 0 | 0.64275716 | 0.90510464 |
| IL1B=IL1R1 | IP-24 | 0 | 0 | 0.7487963 | 0.86673909 |
| IL1B=IL1R2 | IP-24 | 0 | 0 | 0.67869483 | 0.80319579 |
| IL1B=IL1RAP | IP-24 | 0 | 0 | 0.78113242 | 0.92544112 |
| IL1RAP=IL1R1 | IP-24 | 0 | 0 | 0.74260298 | 0.82944239 |
| IL1RN=IL1R1 | IP-24 | 0 | 0 | 0.76440936 | 0.8538446 |
| IL1RN=IL1R2 | IP-24 | 0 | 0 | 0.64550695 | 0.79754795 |
| IL6=IL6R | IP-24 | 0 | 0 | 0.73722553 | 0.84517307 |
| IL6=IL6ST | IP-24 | 0 | 0 | 0.74528097 | 0.83664063 |
| IL6R=IL6ST | IP-24 | 0 | 0 | 0.75758046 | 0.92208143 |
| JAG1=NOTCH: | IP-24 | 0 | 0 | 0.85983576 | 0.97379882 |
| JAG1=NOTCH4 | IP-24 | 0 | 0 | 0.83216059 | 0.90827494 |
| LIFR=IL6ST | IP-24 | 0 | 0 | 0.65444411 | 0.76997549 |
| LPL=LRP1 | IP-24 | 0 | 0 | 0.78912656 | 0.86407203 |
| MST1=MST1R | IP-24 | 0 | 0 | 0.63495104 | 0.7556957 |
| NTS=NTSR1 | IP-24 | 0 | 0 | 0.59105721 | 0.76288558 |
| OSM=IL6ST | IP-24 | 0 | 0 | 0.65706615 | 0.86999335 |
| PGF=FLT1 | IP-24 | 0 | 0 | 0.61593053 | 0.78110821 |
| PLAT=LRP1 | IP-24 | 0 | 0 | 0.50026164 | 0.79655539 |
| PTPRF=INSR | IP-24 | 0 | 0 | 0.66799183 | 0.83277136 |
| SELPLG=SELL | IP-24 | 0 | 0 | 0.71548818 | 0.78113156 |
| SEMA3C=NRP | IP-24 | 0 | 0 | 0.72337821 | 0.90333302 |
| SEMA3C=PLXI | IP-24 | 0 | 0 | 0.83932851 | 0.9794116 |
| SEMA3E=PLXI | IP-24 | 0 | 0 | 0.63991538 | 0.76735837 |
| SEMA4A=PLXI | IP-24 | 0 | 0 | 0.93333921 | 0.99642924 |
| SERPINA1=LRI | IP-24 | 0 | 0 | 0.73957024 | 0.95326918 |
| SHH=PTCH2 | IP-24 | 0 | 0 | 0.57317454 | 0.75711591 |
| TF=TFRC | IP-24 | 0 | 0 | 0.61553629 | 0.79960151 |
| THBS1=CD36 | IP-24 | 0 | 0 | 0.76900079 | 0.86444279 |
| THBS1=LRP1 | IP-24 | 0 | 0 | 0.79761205 | 0.87271309 |
| TIMP1=CD63 | IP-24 | 0 | 0 | 0.83792773 | 0.94487467 |
| TNF=TNFRSF1 | IP-24 | 0 | 0 | 0.8496349 | 0.98307627 |
| TNF=TNFRSF1 | IP-24 | 0 | 0 | 0.84993866 | 0.96027456 |
| TNFSF10=TNF | IP-24 | 0 | 0 | 0.85788875 | 0.95140653 |
| TNFSF10=TNF | IP-24 | 0 | 0 | 0.8872396 | 0.96663646 |
| TNFSF10=TNF | IP-24 | 0 | 0 | 0.83074492 | 0.95786156 |
| TNFSF12=TNF | IP-24 | 0 | 0 | 0.7309688 | 0.85155788 |
| TNFSF14=LTBI | IP-24 | 0 | 0 | 0.77254622 | 0.8454687 |
| VCAN=CD44 | IP-24 | 0 | 0 | 0.66368723 | 0.93158435 |
| VEGFA=FLT1 | IP-24 | 0 | 0 | 0.85888125 | 0.84459775 |
| APOE=LDLR | IP-25 | 0 | 0 | 0.71033301 | 0.82025711 |

| | | | | | |
|---|---|---|---|---|---|
| APOE=LRP1 | IP-25 | 0 | 0 | 0.68992092 | 0.81227406 |
| CCL4=CCR1 | IP-25 | 0 | 0 | 0.52114437 | 0.7884948 |
| CCL4=CCR5 | IP-25 | 0 | 0 | 0.63450093 | 0.80050077 |
| CCL5=CCR1 | IP-25 | 0 | 0 | 0.46258815 | 0.75415386 |
| CD244=CD48 | IP-25 | 0 | 0 | 0.82176415 | 0.84594753 |
| DLL1=NOTCH2 | IP-25 | 0 | 0 | 0.53529571 | 0.69261259 |
| FASLG=FAS | IP-25 | 0 | 0 | 0.59559415 | 0.75534584 |
| GDF11=ACVR2 | IP-25 | 0 | 0 | 0.6101625 | 0.77108246 |
| IFNG=IFNGR1 | IP-25 | 0 | 0 | 0.53994773 | 0.74418423 |
| IFNG=IFNGR2 | IP-25 | 0 | 0 | 0.5218814 | 0.77308264 |
| IL16=CD4 | IP-25 | 0 | 0 | 0.74123794 | 0.87755706 |
| JAG2=NOTCH2 | IP-25 | 0 | 0 | 0.59324368 | 0.73683403 |
| JAG2=NOTCH3 | IP-25 | 0 | 0 | 0.62636228 | 0.76776255 |
| LTA=TNFRSF14 | IP-25 | 0 | 0 | 0.68655561 | 0.82764647 |
| LTA=TNFRSF1A | IP-25 | 0 | 0 | 0.68617078 | 0.8556821 |
| LTA=TNFRSF1 | IP-25 | 0 | 0 | 0.68706715 | 0.83584455 |
| TF=TFRC | IP-25 | 0 | 0 | 0.61553629 | 0.79960151 |
| TGFB1=TGFBR | IP-25 | 0 | 0 | 0.97539308 | 0.85339496 |
| TGFB3=TGFBR | IP-25 | 0 | 0 | 0.65711568 | 0.78011442 |
| TGFBR1=TGFB | IP-25 | 0 | 0 | 0.82071839 | 0.81227659 |
| TNFSF14=TNF | IP-25 | 0 | 0 | 0.81489991 | 0.80667495 |
| TRAF2=TNFRS | IP-25 | 0 | 0 | 0.78187375 | 0.86092716 |
| VEGFB=FLT1 | IP-25 | 0 | 0 | 0.75990897 | 0.80934144 |
| A2M=LRP1 | IP-26 | 0 | 0 | 0.4863614 | 0.73400135 |
| ADAM10=NOT | IP-26 | 0 | 0 | 0.89825815 | 0.90949845 |
| ANXA1=FPR1 | IP-26 | 0 | 0 | 0.72963551 | 0.90819847 |
| ANXA1=FPR2 | IP-26 | 0 | 0 | 0.84316937 | 0.8994962 |
| APP=TNFRSF2 | IP-26 | 0 | 0 | 0.63994307 | 0.82753409 |
| B2M=CD1A | IP-26 | 0 | 0 | 0.7125452 | 0.87306562 |
| BMPR2=ACVR | IP-26 | 0 | 0 | 0.92206326 | 0.9552782 |
| CALR=LRP1 | IP-26 | 0 | 0 | 0.58778819 | 0.81440941 |
| CCL2=CCR5 | IP-26 | 0 | 0 | 0.7719236 | 0.84308466 |
| CCL28=CCR3 | IP-26 | 0 | 0 | 0.51852326 | 0.77709353 |
| CCL3=CCR5 | IP-26 | 0 | 0 | 0.80923394 | 0.87668448 |
| CCL3L3=CCR5 | IP-26 | 0 | 0 | 0.81800447 | 0.87444089 |
| CCL4=CCR1 | IP-26 | 0 | 0 | 0.52114437 | 0.7884948 |
| CCL5=CCR3 | IP-26 | 0 | 0 | 0.50505271 | 0.7615359 |
| CD244=CD48 | IP-26 | 0 | 0 | 0.82176415 | 0.84594753 |
| CD47=SIRPA | IP-26 | 0 | 0 | 0.64990708 | 0.8714872 |
| CD6=ALCAM | IP-26 | 0 | 0 | 0.52863435 | 0.81077491 |
| DLL1=NOTCH3 | IP-26 | 0 | 0 | 0.59040003 | 0.75106466 |
| EFNA1=EPHB1 | IP-26 | 0 | 0 | 0.57398927 | 0.76229272 |
| EFNA4=EPHA2 | IP-26 | 0 | 0 | 0.60484267 | 0.8022388 |
| EFNA5=EPHA2 | IP-26 | 0 | 0 | 0.60478794 | 0.7883101 |
| EFNA5=EPHB2 | IP-26 | 0 | 0 | 0.71505684 | 0.83868372 |
| EFNB1=EPHB2 | IP-26 | 0 | 0 | 0.86682713 | 0.88139107 |
| EFNB1=EPHB3 | IP-26 | 0 | 0 | 0.77904273 | 0.85073096 |
| F2=F2RL1 | IP-26 | 0 | 0 | 0.58468516 | 0.77433865 |
| F2=THBD | IP-26 | 0 | 0 | 0.59527841 | 0.78364551 |
| FGFR1=CDH1 | IP-26 | 0 | 0 | 0.57267629 | 0.76230883 |

| | | | | | |
|---|---|---|---|---|---|
| GAS6=AXL | IP-26 | 0 | 0 | 0.77454072 | 0.81100794 |
| HLA-A=LILRB1 | IP-26 | 0 | 0 | 0.85560044 | 0.86901743 |
| HLA-A=LILRB2 | IP-26 | 0 | 0 | 0.67962281 | 0.87146779 |
| HLA-B=LILRB2 | IP-26 | 0 | 0 | 0.70767507 | 0.88257716 |
| HLA-G=LILRB1 | IP-26 | 0 | 0 | 0.92025046 | 0.88534952 |
| HLA-G=LILRB2 | IP-26 | 0 | 0 | 0.80611218 | 0.88734373 |
| IFNG=IFNGR2 | IP-26 | 0 | 0 | 0.5218814 | 0.77308264 |
| IGF2=INSR | IP-26 | 0 | 0 | 0.60109031 | 0.79364999 |
| IL16=CD4 | IP-26 | 0 | 0 | 0.74123794 | 0.87755706 |
| INSL3=RXFP2 | IP-26 | 0 | 0 | 0.68462865 | 0.78000037 |
| ITGAL=ICAM1 | IP-26 | 0 | 0 | 0.87294769 | 0.86610437 |
| JAG1=NOTCH3 | IP-26 | 0 | 0 | 0.75816003 | 0.84540297 |
| LTA=LTBR | IP-26 | 0 | 0 | 0.66879107 | 0.8509929 |
| PTPRF=INSR | IP-26 | 0 | 0 | 0.66799183 | 0.83277136 |
| RSPO1=LGR4 | IP-26 | 0 | 0 | 0.55037624 | 0.76861457 |
| SELPLG=SELL | IP-26 | 0 | 0 | 0.71548818 | 0.78113156 |
| SEMA3C=NRP | IP-26 | 0 | 0 | 0.72337821 | 0.90333302 |
| SEMA4D=PLXI | IP-26 | 0 | 0 | 0.73743573 | 0.85482063 |
| SEMA7A=PLXI | IP-26 | 0 | 0 | 0.65993156 | 0.82980314 |
| SERPINC1=LRI | IP-26 | 0 | 0 | 0.52842581 | 0.77055291 |
| TGFB1=ACVRL | IP-26 | 0 | 0 | 0.79368843 | 0.85170966 |
| TGFB1=APP | IP-26 | 0 | 0 | 0.83918292 | 0.90158678 |
| TGFB1=SDC2 | IP-26 | 0 | 0 | 0.79230495 | 0.87143661 |
| TGFB3=ACVRL | IP-26 | 0 | 0 | 0.60363713 | 0.80558319 |
| TNF=TNFRSF1 | IP-26 | 0 | 0 | 0.8496349 | 0.98307627 |
| TNF=TNFRSF1 | IP-26 | 0 | 0 | 0.84993866 | 0.96027456 |
| TNFSF12=TNF | IP-26 | 0 | 0 | 0.7309688 | 0.85155788 |
| TNFSF14=LTBI | IP-26 | 0 | 0 | 0.77254622 | 0.8454687 |
| TRAF2=TNFRS | IP-26 | 0 | 0 | 0.78187375 | 0.86092716 |
| VEGFA=NRP1 | IP-26 | 0 | 0 | 0.67191177 | 0.84496415 |
| VEGFB=NRP1 | IP-26 | 0 | 0 | 0.63766862 | 0.81379153 |
| XCL1=XCR1 | IP-26 | 0 | 0 | 0.56205717 | 0.75659613 |
| CCL5=CCR5 | IP-27 | 0 | 0 | 0.60099082 | 0.77927246 |
| DLL1=NOTCH1 | IP-27 | 0 | 0 | 0.59064493 | 0.68955635 |
| DLL1=NOTCH2 | IP-27 | 0 | 0 | 0.53529571 | 0.69261259 |
| F2=F2RL1 | IP-27 | 0 | 0 | 0.58468516 | 0.77433865 |
| F2=THBD | IP-27 | 0 | 0 | 0.59527841 | 0.78364551 |
| FAT4=DCHS1 | IP-27 | 0 | 0 | 0.55130249 | 0.72850405 |
| GDF11=ACVR2 | IP-27 | 0 | 0 | 0.6101625 | 0.77108246 |
| IGF2=IGF1R | IP-27 | 0 | 0 | 0.59468691 | 0.78257292 |
| IGF2=IGF2R | IP-27 | 0 | 0 | 0.57535173 | 0.76377911 |
| IL5=CSF2RB | IP-27 | 0 | 0 | 0.60557309 | 0.75817301 |
| JAG2=NOTCH1 | IP-27 | 0 | 0 | 0.64014498 | 0.726604 |
| LAMA2=RPSA | IP-27 | 0 | 0 | 0.55004722 | 0.72923306 |
| TGFB3=TGFBF | IP-27 | 0 | 0 | 0.6505711 | 0.76998295 |
| VEGFB=NRP1 | IP-27 | 0 | 0 | 0.63766862 | 0.81379153 |
| WNT1=RYK | IP-27 | 0 | 0 | 0.68481992 | 0.82054329 |
| WNT3=FZD8 | IP-27 | 0 | 0 | 0.56833056 | 0.75804153 |
| CCL13=ACKR4 | IP-28 | 0.1002 | 0.0237 | 0.47097408 | 0.00457645 |
| CD274=PDCD1 | IP-28 | 0.1002 | 0.0237 | 0.55027644 | 0.7543037 |

| | | | | | |
|---|---|---|---|---|---|
| CXCL10=CXCR | IP-28 | 0.1002 | 0.0237 | 0.61989852 | 0.78505436 |
| CXCL11=CXCR | IP-28 | 0.1002 | 0.0237 | 0.59616263 | 0.771329 |
| CXCL9=CXCR3 | IP-28 | 0.1002 | 0.0237 | 0.5895112 | 0.78577405 |
| EFNA4=EPHA( | IP-28 | 0.1002 | 0.0237 | 0.56327182 | 0.7892322 |
| EFNB2=EPHB1 | IP-28 | 0.1002 | 0.0237 | 0.55738138 | 0.80852037 |
| IL17B=IL17RB | IP-28 | 0.1002 | 0.0237 | 0.51025689 | 0.78567141 |
| IL19=IL20RB | IP-28 | 0.1002 | 0.0237 | 0.50182437 | 0.76799995 |
| IL22=IL10RB | IP-28 | 0.1002 | 0.0237 | 0.60312174 | 0.76309067 |
| PDCD1LG2=PI | IP-28 | 0.1002 | 0.0237 | 0.5679087 | 0.76458293 |
| TNFSF13=TNF | IP-28 | 0.1002 | 0.0237 | 0.56872965 | 0.76516172 |
| TNFSF13B=TN | IP-28 | 0.1002 | 0.0237 | 0.57316522 | 0.76560769 |
| TNFSF13B=TN | IP-28 | 0.1002 | 0.0237 | 0.51030319 | 0.77337441 |
| WNT3A=FZD6 | IP-28 | 0.1002 | 0.0237 | 0.56222527 | 0.78315864 |
| CXCL1=CXCR1 | IP-29 | 0 | 0 | 0.64009773 | 0.81732317 |
| CXCL1=CXCR2 | IP-29 | 0 | 0 | 0.63827843 | 0.81465275 |
| CXCL2=CXCR1 | IP-29 | 0 | 0 | 0.70532745 | 0.85440808 |
| CXCL2=CXCR2 | IP-29 | 0 | 0 | 0.69515763 | 0.85008323 |
| CXCL3=CXCR1 | IP-29 | 0 | 0 | 0.690754 | 0.85318129 |
| CXCL3=CXCR2 | IP-29 | 0 | 0 | 0.67447738 | 0.84962529 |
| CXCL5=CXCR1 | IP-29 | 0 | 0 | 0.61226566 | 0.78233283 |
| CXCL5=CXCR2 | IP-29 | 0 | 0 | 0.61566311 | 0.78293309 |
| CXCL6=CXCR1 | IP-29 | 0 | 0 | 0.5614179 | 0.78115438 |
| CXCL6=CXCR2 | IP-29 | 0 | 0 | 0.56205436 | 0.78043795 |
| EFNB2=EPHB4 | IP-29 | 0 | 0 | 0.52599006 | 0.82817489 |
| FGF22=FGFR2 | IP-29 | 0 | 0 | 0.5098951 | 0.76764253 |
| FGF23=FGFR2 | IP-29 | 0 | 0 | 0.54282509 | 0.7663766 |
| FGF5=FGFR2 | IP-29 | 0 | 0 | 0.54723963 | 0.76440621 |
| IL1A=IL1R1 | IP-29 | 0 | 0 | 0.63163394 | 0.85474413 |
| IL1B=IL1R1 | IP-29 | 0 | 0 | 0.7487963 | 0.86673909 |
| IL1RN=IL1R1 | IP-29 | 0 | 0 | 0.76440936 | 0.8538446 |
| MMP9=CD44 | IP-29 | 0 | 0 | 0.59510155 | 0.77767395 |
| PPBP=CXCR1 | IP-29 | 0 | 0 | 0.59429064 | 0.77060198 |
| PPBP=CXCR2 | IP-29 | 0 | 0 | 0.5904397 | 0.76913702 |
| SPP1=CD44 | IP-29 | 0 | 0 | 0.6266785 | 0.76294716 |
| CD28=CD86 | IP-3 | 0.0065 | 0.008 | 0.54550868 | 0.81234109 |
| CD40LG=CD4( | IP-3 | 0.0065 | 0.008 | 0.58082821 | 0.80293737 |
| FLT3LG=FLT3 | IP-3 | 0.0065 | 0.008 | 0.63756932 | 0.77772317 |
| SEMA3A=NRP | IP-3 | 0.0065 | 0.008 | 0.56218263 | 0.78071232 |
| WNT7A=FZD5 | IP-3 | 0.0065 | 0.008 | 0.52313987 | 0.76637496 |
| CCL13=CCR3 | IP-30 | 0.0082 | 0.0028 | 0.22818025 | 0.00714358 |
| CCL8=CCR5 | IP-30 | 0.0082 | 0.0028 | 0.66466171 | 0.78653071 |
| CSF2=CSF2RA | IP-30 | 0.0082 | 0.0028 | 0.62954724 | 0.67469353 |
| CSF2=CSF2RB | IP-30 | 0.0082 | 0.0028 | 0.60788543 | 0.66806116 |
| CSF2=CSF3R | IP-30 | 0.0082 | 0.0028 | 0.60942533 | 0.66890786 |
| EFNB1=EPHB1 | IP-30 | 0.0082 | 0.0028 | 0.67504948 | 0.8248148 |
| GAS6=AXL | IP-30 | 0.0082 | 0.0028 | 0.77454072 | 0.81100794 |
| IL22=IL10RB | IP-30 | 0.0082 | 0.0028 | 0.60312174 | 0.76309067 |
| INSR=CEACAN | IP-30 | 0.0082 | 0.0028 | 0.63390094 | 0.77050135 |
| NRG1=ERBB3 | IP-30 | 0.0082 | 0.0028 | 0.57637479 | 0.81914395 |
| SEMA3E=PLXN | IP-30 | 0.0082 | 0.0028 | 0.63991538 | 0.76735837 |

| | | | | | |
|---|---|---|---|---|---|
| TAC1=TACR2 | IP-30 | 0.0082 | 0.0028 | 0.54941368 | 0.76574859 |
| WNT3A=FZD2 | IP-30 | 0.0082 | 0.0028 | 0.64991298 | 0.8219015 |
| WNT3A=FZD5 | IP-30 | 0.0082 | 0.0028 | 0.65001846 | 0.79062061 |
| WNT3A=FZD7 | IP-30 | 0.0082 | 0.0028 | 0.53400732 | 0.80430691 |
| WNT3A=RYK | IP-30 | 0.0082 | 0.0028 | 0.65901372 | 0.83042884 |
| ADAM9=ITGA | IP-31 | 0 | 0 | 0.83387244 | 0.95552805 |
| CALCA=CALCR | IP-31 | 0 | 0 | 0.5542414 | 0.78168041 |
| CD28=CD86 | IP-31 | 0 | 0 | 0.54550868 | 0.81234109 |
| CD40LG=CD4( | IP-31 | 0 | 0 | 0.58082821 | 0.80293737 |
| CD86=CD28 | IP-31 | 0 | 0 | 0.55096425 | 0.81566693 |
| CD86=CTLA4 | IP-31 | 0 | 0 | 0.5220613 | 0.79955309 |
| CRH=CRHR1 | IP-31 | 0 | 0 | 0.50075688 | 0.01402565 |
| CSF1=CSF1R | IP-31 | 0 | 0 | 0.75004398 | 0.81006548 |
| CTLA4=CD86 | IP-31 | 0 | 0 | 0.51559262 | 0.78531206 |
| DKK1=KREME | IP-31 | 0 | 0 | 0.58533705 | 0.64402008 |
| EPO=EPOR | IP-31 | 0 | 0 | 0.61688504 | 0.77389236 |
| FBN1=ITGAV | IP-31 | 0 | 0 | 0.5780073 | 0.80742082 |
| FBN1=ITGB1 | IP-31 | 0 | 0 | 0.57209971 | 0.79260136 |
| FN1=ITGA4 | IP-31 | 0 | 0 | 0.78351002 | 0.82671942 |
| FN1=ITGA5 | IP-31 | 0 | 0 | 0.78682058 | 0.84208529 |
| FN1=ITGAV | IP-31 | 0 | 0 | 0.81417799 | 0.84489418 |
| FN1=ITGB1 | IP-31 | 0 | 0 | 0.74769308 | 0.81342652 |
| ICAM1=ITGAL | IP-31 | 0 | 0 | 0.99784278 | 0.92810535 |
| ICOS=ICOSLG | IP-31 | 0 | 0 | 0.55299903 | 0.80123717 |
| ICOSLG=ICOS | IP-31 | 0 | 0 | 0.5641786 | 0.81214186 |
| IL10=IL10RA | IP-31 | 0 | 0 | 0.89545338 | 0.86259435 |
| IL10=IL10RB | IP-31 | 0 | 0 | 0.87909324 | 0.88338511 |
| IL27=IL27RA | IP-31 | 0 | 0 | 0.94073811 | 0.92285822 |
| INHBB=ACVR1 | IP-31 | 0 | 0 | 0.71683626 | 0.74999524 |
| INHBB=ACVR2 | IP-31 | 0 | 0 | 0.69419724 | 0.74336803 |
| INSL3=RXFP2 | IP-31 | 0 | 0 | 0.68462865 | 0.78000037 |
| INSR=ADRB2 | IP-31 | 0 | 0 | 0.93664696 | 0.94015887 |
| JAG1=NOTCH | IP-31 | 0 | 0 | 0.87470324 | 0.92436765 |
| LGALS9=HAV( | IP-31 | 0 | 0 | 0.93860412 | 0.87878614 |
| MMP9=CD44 | IP-31 | 0 | 0 | 0.59510155 | 0.77767395 |
| NRG1=ERBB3 | IP-31 | 0 | 0 | 0.57637479 | 0.81914395 |
| PCSK9=LDLR | IP-31 | 0 | 0 | 0.69335189 | 0.77340033 |
| PRL=PRLR | IP-31 | 0 | 0 | 0.60118138 | 0.7617163 |
| SEMA3C=PLXI | IP-31 | 0 | 0 | 0.83932851 | 0.9794116 |
| SIRPA=CD47 | IP-31 | 0 | 0 | 0.75251255 | 0.90748531 |
| SPP1=CD44 | IP-31 | 0 | 0 | 0.6266785 | 0.76294716 |
| THBS1=CD47 | IP-31 | 0 | 0 | 0.82010142 | 0.84499401 |
| TNFSF15=TNF | IP-31 | 0 | 0 | 0.58401156 | 0.80751455 |
| WNT3A=FZD8 | IP-31 | 0 | 0 | 0.57805252 | 0.79742624 |
| ADAM17=NO | IP-32 | 0 | 0 | 0.93244589 | 0.91031936 |
| ALCAM=CD6 | IP-32 | 0 | 0 | 0.59957352 | 0.84002104 |
| APOE=VLDLR | IP-32 | 0 | 0 | 0.61285988 | 0.77908389 |
| BAG6=NCR3 | IP-32 | 0 | 0 | 0.52679041 | 0.65955997 |
| BMP7=ACVR1 | IP-32 | 0 | 0 | 0.62387375 | 0.82337927 |
| BMP7=ACVR2 | IP-32 | 0 | 0 | 0.62577649 | 0.80486923 |

| | | | | | |
|---|---|---|---|---|---|
| BMP7=ACVR2 | IP-32 | 0 | 0 | 0.61061404 | 0.80755253 |
| BMP7=BMPR1 | IP-32 | 0 | 0 | 0.57728798 | 0.78136696 |
| BMP7=BMPR1 | IP-32 | 0 | 0 | 0.60142136 | 0.79542963 |
| BMP7=BMPR2 | IP-32 | 0 | 0 | 0.6123408 | 0.80945918 |
| BMPR2=BMPI | IP-32 | 0 | 0 | 0.75794761 | 0.83663718 |
| CALCB=RAMP | IP-32 | 0 | 0 | 0.54482358 | 0.75821038 |
| CD48=CD244 | IP-32 | 0 | 0 | 0.86741813 | 0.86376962 |
| CD58=CD2 | IP-32 | 0 | 0 | 0.62254549 | 0.77141118 |
| CDH1=PTPRM | IP-32 | 0 | 0 | 0.59283266 | 0.79693049 |
| CXCL16=CXCR | IP-32 | 0 | 0 | 0.62385603 | 0.85146955 |
| EFNA4=EPHA4 | IP-32 | 0 | 0 | 0.61484688 | 0.80233147 |
| EFNA5=EPHA4 | IP-32 | 0 | 0 | 0.60571641 | 0.75562706 |
| EFNB1=EPHB3 | IP-32 | 0 | 0 | 0.77904273 | 0.85073096 |
| EFNB2=EPHA4 | IP-32 | 0 | 0 | 0.54559921 | 0.79608261 |
| EFNB2=EPHB2 | IP-32 | 0 | 0 | 0.61831418 | 0.85337707 |
| EFNB2=EPHB3 | IP-32 | 0 | 0 | 0.59015387 | 0.82928359 |
| EPHB2=EFNB2 | IP-32 | 0 | 0 | 0.55180981 | 0.82865093 |
| F12=CD93 | IP-32 | 0 | 0 | 0.73361191 | 0.8310069 |
| GAS6=MERTK | IP-32 | 0 | 0 | 0.83377044 | 0.85036214 |
| HLA-A=KIR3DI | IP-32 | 0 | 0 | 0.5168139 | 0.64416742 |
| HLA-B=KIR3DI | IP-32 | 0 | 0 | 0.53633565 | 0.64819374 |
| HLA-C=KIR2DI | IP-32 | 0 | 0 | 0.53578013 | 0.65594848 |
| HLA-C=KIR2DI | IP-32 | 0 | 0 | 0.53392319 | 0.64432278 |
| HLA-E=KLRC2 | IP-32 | 0 | 0 | 0.5071642 | 0.63253053 |
| HLA-E=KLRD1 | IP-32 | 0 | 0 | 0.51682783 | 0.64103884 |
| ICAM3=ITGAL | IP-32 | 0 | 0 | 0.6774393 | 0.74243302 |
| IHH=PTCH1 | IP-32 | 0 | 0 | 0.5676963 | 0.76322693 |
| IL15=IL2RB | IP-32 | 0 | 0 | 0.56477991 | 0.76431507 |
| IL15=IL2RG | IP-32 | 0 | 0 | 0.7447103 | 0.83650527 |
| IL18=IL18R1 | IP-32 | 0 | 0 | 0.7049463 | 0.7955952 |
| IL4=IL13RA1 | IP-32 | 0 | 0 | 0.64355142 | 0.82001589 |
| IL4=IL2RG | IP-32 | 0 | 0 | 0.60173746 | 0.78170144 |
| IL4=IL4R | IP-32 | 0 | 0 | 0.61612416 | 0.79190901 |
| IL4R=IL2RG | IP-32 | 0 | 0 | 0.68262718 | 0.77525508 |
| IL6=IL6R | IP-32 | 0 | 0 | 0.73722553 | 0.84517307 |
| IL6=IL6ST | IP-32 | 0 | 0 | 0.74528097 | 0.83664063 |
| IL7=IL2RG | IP-32 | 0 | 0 | 0.59861506 | 0.76789009 |
| ITGB1BP1=ITG | IP-32 | 0 | 0 | 0.77062634 | 0.79348204 |
| JAG1=NOTCH1 | IP-32 | 0 | 0 | 0.75816003 | 0.84540297 |
| LIF=IL6ST | IP-32 | 0 | 0 | 0.63159173 | 0.8480836 |
| MADCAM1=IT | IP-32 | 0 | 0 | 0.53061945 | 0.79727471 |
| MADCAM1=IT | IP-32 | 0 | 0 | 0.52320922 | 0.77296094 |
| MICA=KLRK1 | IP-32 | 0 | 0 | 0.61936661 | 0.74477466 |
| NCR3LG1=NCI | IP-32 | 0 | 0 | 0.55378099 | 0.69972482 |
| NOTCH1=DLL1 | IP-32 | 0 | 0 | 0.59422226 | 0.6980767 |
| NRG1=ERBB2 | IP-32 | 0 | 0 | 0.5466478 | 0.75337675 |
| PDGFD=PDGF | IP-32 | 0 | 0 | 0.50020011 | 0.63150855 |
| PLAU=PLAUR | IP-32 | 0 | 0 | 0.6307255 | 0.84984968 |
| POMC=MC1R | IP-32 | 0 | 0 | 0.62361958 | 0.74033156 |
| PTPRJ=PDGFR | IP-32 | 0 | 0 | 0.56967198 | 0.6807627 |

| | | | | | |
|---|---|---|---|---|---|
| PVR=CD226 | IP-32 | 0 | 0 | 0.60952202 | 0.8080848 |
| RARRES2=CM | IP-32 | 0 | 0 | 0.58790506 | 0.79906184 |
| RELN=LRP8 | IP-32 | 0 | 0 | 0.55563125 | 0.78813104 |
| RELN=VLDLR | IP-32 | 0 | 0 | 0.53941563 | 0.77327668 |
| SHH=PTCH1 | IP-32 | 0 | 0 | 0.55861069 | 0.74641541 |
| TGFB1=TGFBF | IP-32 | 0 | 0 | 0.81054739 | 0.8012184 |
| TGFB1=TGFBF | IP-32 | 0 | 0 | 0.97539308 | 0.85339496 |
| TGFB1=TGFBF | IP-32 | 0 | 0 | 0.54219406 | 0.68103431 |
| TGFB2=TGFBF | IP-32 | 0 | 0 | 0.60926063 | 0.79351529 |
| TGFB2=TGFBF | IP-32 | 0 | 0 | 0.60863144 | 0.79644552 |
| TGFB2=TGFBF | IP-32 | 0 | 0 | 0.54320133 | 0.77114725 |
| TGFB3=TGFBF | IP-32 | 0 | 0 | 0.6505711 | 0.76998295 |
| ULBP2=KLRK1 | IP-32 | 0 | 0 | 0.52977936 | 0.7507739 |
| VEGFA=FLT1 | IP-32 | 0 | 0 | 0.85888125 | 0.84459775 |
| BMP4=BMPR1 | IP-4 | 0 | 0.0022 | 0.61344759 | 0.78911158 |
| BMP4=BMPR1 | IP-4 | 0 | 0.0022 | 0.646575 | 0.80084393 |
| BMP4=BMPR2 | IP-4 | 0 | 0.0022 | 0.69038242 | 0.81661979 |
| BMP6=BMPR1 | IP-4 | 0 | 0.0022 | 0.59114087 | 0.75839481 |
| BMP6=BMPR1 | IP-4 | 0 | 0.0022 | 0.61354369 | 0.76322688 |
| BMP6=BMPR2 | IP-4 | 0 | 0.0022 | 0.64830978 | 0.77737147 |
| CCL8=CCR5 | IP-4 | 0 | 0.0022 | 0.66466171 | 0.78653071 |
| CDH1=ITGAE | IP-4 | 0 | 0.0022 | 0.6434374 | 0.8502289 |
| CDH1=PTPRM | IP-4 | 0 | 0.0022 | 0.59283266 | 0.79693049 |
| CX3CL1=CX3C | IP-4 | 0 | 0.0022 | 0.56985381 | 0.74789456 |
| DLK1=NOTCH1 | IP-4 | 0 | 0.0022 | 0.5679145 | 0.75366957 |
| EGF=ERBB2 | IP-4 | 0 | 0.0022 | 0.56859923 | 0.76211995 |
| FBN1=ITGAV | IP-4 | 0 | 0.0022 | 0.5780073 | 0.80742082 |
| FBN1=ITGB1 | IP-4 | 0 | 0.0022 | 0.57209971 | 0.79260136 |
| FGF2=FGFR1 | IP-4 | 0 | 0.0022 | 0.58397679 | 0.73507273 |
| FGF2=FGFRL1 | IP-4 | 0 | 0.0022 | 0.60762262 | 0.75308915 |
| FGF2=SDC2 | IP-4 | 0 | 0.0022 | 0.59713788 | 0.75958532 |
| FGF2=SDC3 | IP-4 | 0 | 0.0022 | 0.63108776 | 0.76547981 |
| FGF2=SDC4 | IP-4 | 0 | 0.0022 | 0.61810207 | 0.75318751 |
| FGF23=FGFR1 | IP-4 | 0 | 0.0022 | 0.58199269 | 0.75651258 |
| IHH=PTCH1 | IP-4 | 0 | 0.0022 | 0.5676963 | 0.76322693 |
| IL4=IL13RA1 | IP-4 | 0 | 0.0022 | 0.64355142 | 0.82001589 |
| IL4=IL2RG | IP-4 | 0 | 0.0022 | 0.60173746 | 0.78170144 |
| IL4=IL4R | IP-4 | 0 | 0.0022 | 0.61612416 | 0.79190901 |
| PDGFB=PDGFI | IP-4 | 0 | 0.0022 | 0.5621465 | 0.75688846 |
| RSPO1=LGR6 | IP-4 | 0 | 0.0022 | 0.54466832 | 0.73696215 |
| SELP=SELPLG | IP-4 | 0 | 0.0022 | 0.64869602 | 0.77210416 |
| TGFB2=TGFBF | IP-4 | 0 | 0.0022 | 0.60926063 | 0.79351529 |
| TGFB2=TGFBF | IP-4 | 0 | 0.0022 | 0.60863144 | 0.79644552 |
| TGFB2=TGFBF | IP-4 | 0 | 0.0022 | 0.54320133 | 0.77114725 |
| TNFSF18=TNF | IP-4 | 0 | 0.0022 | 0.54130329 | 0.7559906 |
| WNT4=FZD6 | IP-4 | 0 | 0.0022 | 0.57441114 | 0.76226496 |
| APP=TNFRSF2 | IP-5 | 0 | 0.0005 | 0.63994307 | 0.82753409 |
| EFNA1=EPHA2 | IP-5 | 0 | 0.0005 | 0.5427516 | 0.75759198 |
| EFNA4=EPHA2 | IP-5 | 0 | 0.0005 | 0.60484267 | 0.8022388 |
| EFNA5=EPHA2 | IP-5 | 0 | 0.0005 | 0.60478794 | 0.7883101 |

| | | | | | |
|---|---|---|---|---|---|
| FGFR1=CDH1 | IP-5 | 0 | 0.0005 | 0.57267629 | 0.76230883 |
| VEGFA=NRP1 | IP-5 | 0 | 0.0005 | 0.67191177 | 0.84496415 |
| EPHB2=EFNB2 | IP-6 | 0.0325 | 0.0637 | 0.55180981 | 0.82865093 |
| F12=GP1BA | IP-6 | 0.0325 | 0.0637 | 0.57765826 | 0.75142559 |
| IL1A=IL1R1 | IP-6 | 0.0325 | 0.0637 | 0.63163394 | 0.85474413 |
| IL1A=IL1R2 | IP-6 | 0.0325 | 0.0637 | 0.59947556 | 0.7978409 |
| IL1A=IL1RAP | IP-6 | 0.0325 | 0.0637 | 0.64275716 | 0.90510464 |
| SELPLG=SELP | IP-6 | 0.0325 | 0.0637 | 0.5701251 | 0.75664777 |
| A2M=LRP1 | IP-7 | 0 | 0 | 0.4863614 | 0.73400135 |
| CD6=ALCAM | IP-7 | 0 | 0 | 0.52863435 | 0.81077491 |
| JAG2=NOTCH3 | IP-7 | 0 | 0 | 0.62636228 | 0.76776255 |
| LTA=LTBR | IP-7 | 0 | 0 | 0.66879107 | 0.8509929 |
| LTA=TNFRSF14 | IP-7 | 0 | 0 | 0.68655561 | 0.82764647 |
| LTA=TNFRSF1A | IP-7 | 0 | 0 | 0.68617078 | 0.8556821 |
| LTA=TNFRSF1 | IP-7 | 0 | 0 | 0.68706715 | 0.83584455 |
| PLG=F2RL1 | IP-7 | 0 | 0 | 0.71049921 | 0.81021998 |
| SEMA4D=PLXI | IP-7 | 0 | 0 | 0.71390705 | 0.7915268 |
| TGFB3=ACVRL | IP-7 | 0 | 0 | 0.60363713 | 0.80558319 |
| APOE=LDLR | IP-8 | 0 | 0.0005 | 0.71033301 | 0.82025711 |
| APOE=LRP1 | IP-8 | 0 | 0.0005 | 0.68992092 | 0.81227406 |
| APOE=LRP8 | IP-8 | 0 | 0.0005 | 0.70059912 | 0.79911382 |
| BGN=TLR2 | IP-8 | 0 | 0.0005 | 0.61274931 | 0.77720905 |
| BGN=TLR4 | IP-8 | 0 | 0.0005 | 0.60692989 | 0.77783889 |
| CALCB=CALCR | IP-8 | 0 | 0.0005 | 0.62981143 | 0.79077866 |
| CCL16=CCR1 | IP-8 | 0 | 0.0005 | 0.59733141 | 0.75165882 |
| CCL16=CCR2 | IP-8 | 0 | 0.0005 | 0.58781903 | 0.74950182 |
| CCL23=CCR1 | IP-8 | 0 | 0.0005 | 0.65431511 | 0.76377941 |
| CCL8=CCR1 | IP-8 | 0 | 0.0005 | 0.71362018 | 0.80929468 |
| CCL8=CCR2 | IP-8 | 0 | 0.0005 | 0.70129007 | 0.8041171 |
| EDN3=EDNRB | IP-8 | 0 | 0.0005 | 0.63156584 | 0.75858673 |
| EFNA5=EPHB2 | IP-8 | 0 | 0.0005 | 0.71505684 | 0.83868372 |
| EFNB1=EPHB1 | IP-8 | 0 | 0.0005 | 0.67504948 | 0.8248148 |
| F9=LRP1 | IP-8 | 0 | 0.0005 | 0.59657006 | 0.69133432 |
| ICAM3=ITGB2 | IP-8 | 0 | 0.0005 | 0.64164351 | 0.83250858 |
| OSM=IL6ST | IP-8 | 0 | 0.0005 | 0.65706615 | 0.86999335 |
| TGFB1=SDC2 | IP-8 | 0 | 0.0005 | 0.79230495 | 0.87143661 |
| CALR=ITGA3 | IP-9 | 0.0012 | 0.9662 | 0.56536868 | 0.75925729 |
| DLL4=NOTCH1 | IP-9 | 0.0012 | 0.9662 | 0.68094361 | 0.76623646 |
| DLL4=NOTCH2 | IP-9 | 0.0012 | 0.9662 | 0.67875239 | 0.77331019 |
| DLL4=NOTCH3 | IP-9 | 0.0012 | 0.9662 | 0.63692066 | 0.76085038 |
| DLL4=NOTCH4 | IP-9 | 0.0012 | 0.9662 | 0.66357591 | 0.76968784 |
| GDF5=ACVR2I | IP-9 | 0.0012 | 0.9662 | 0.57304632 | 0.77233837 |
| GDF5=BMPR1 | IP-9 | 0.0012 | 0.9662 | 0.55352967 | 0.76157788 |
| GDF5=BMPR1 | IP-9 | 0.0012 | 0.9662 | 0.56529923 | 0.76602605 |
| GDF5=BMPR2 | IP-9 | 0.0012 | 0.9662 | 0.56662413 | 0.76786315 |
| GDF9=BMPR1 | IP-9 | 0.0012 | 0.9662 | 0.57152717 | 0.78369717 |
| IL11=IL11RA | IP-9 | 0.0012 | 0.9662 | 0.5778172 | 0.74621738 |
| IL11=IL6ST | IP-9 | 0.0012 | 0.9662 | 0.59191561 | 0.74994687 |
| INHA=TGFBR3 | IP-9 | 0.0012 | 0.9662 | 0.540287 | 0.71692714 |
| LHB=LHCGR | IP-9 | 0.0012 | 0.9662 | 0.5986825 | 0.71126408 |

| | | | | |
|---|---|---|---|---|
| NODAL=ACVR IP-9 | 0.0012 | 0.9662 | 0.5812556 | 0.79199569 |
| RARRES2=CM IP-9 | 0.0012 | 0.9662 | 0.58790506 | 0.79906184 |
| TIMP2=ITGA3 IP-9 | 0.0012 | 0.9662 | 0.57981617 | 0.79290749 |
