## Supplementary Figures for "NK cell-monocyte crosstalk underlies NK cell activation in severe COVID-19"

### SUPPLEMENTARY MATERIALS

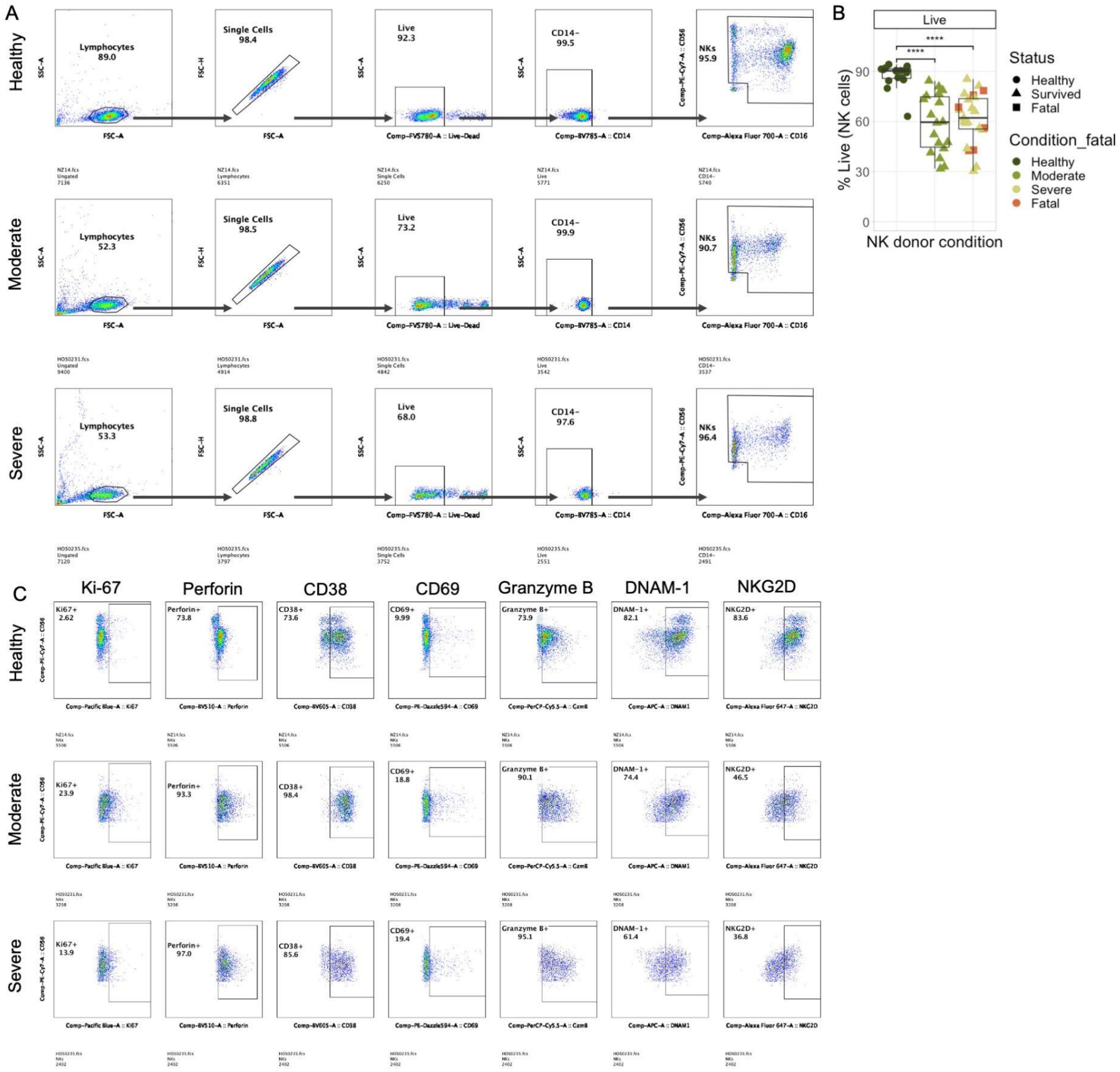


**Supplementary Figure 1: Gating strategy and representative flow plots for primary NK cell phenotyping experiments.** A) Representative flow plots showing the gating strategy used to identify NK cells in NK cell phenotyping experiments. Example plots are shown for a healthy donor (top), moderate COVID-19 donor (middle), and severe COVID-19 donor (bottom). B) Boxplot showing the percentage of NK cells that were live (negative for eFluor 780 fixable viability dye) in each sample. Statistical significance values were determined using a Wilcoxon rank sum test. C) Representative flow plots showing the expression of all phenotypic markers quantified in main text Fig. 2. Example plots are shown for a healthy donor (top), moderate COVID-19 donor (middle), and severe COVID-19 donor (bottom).

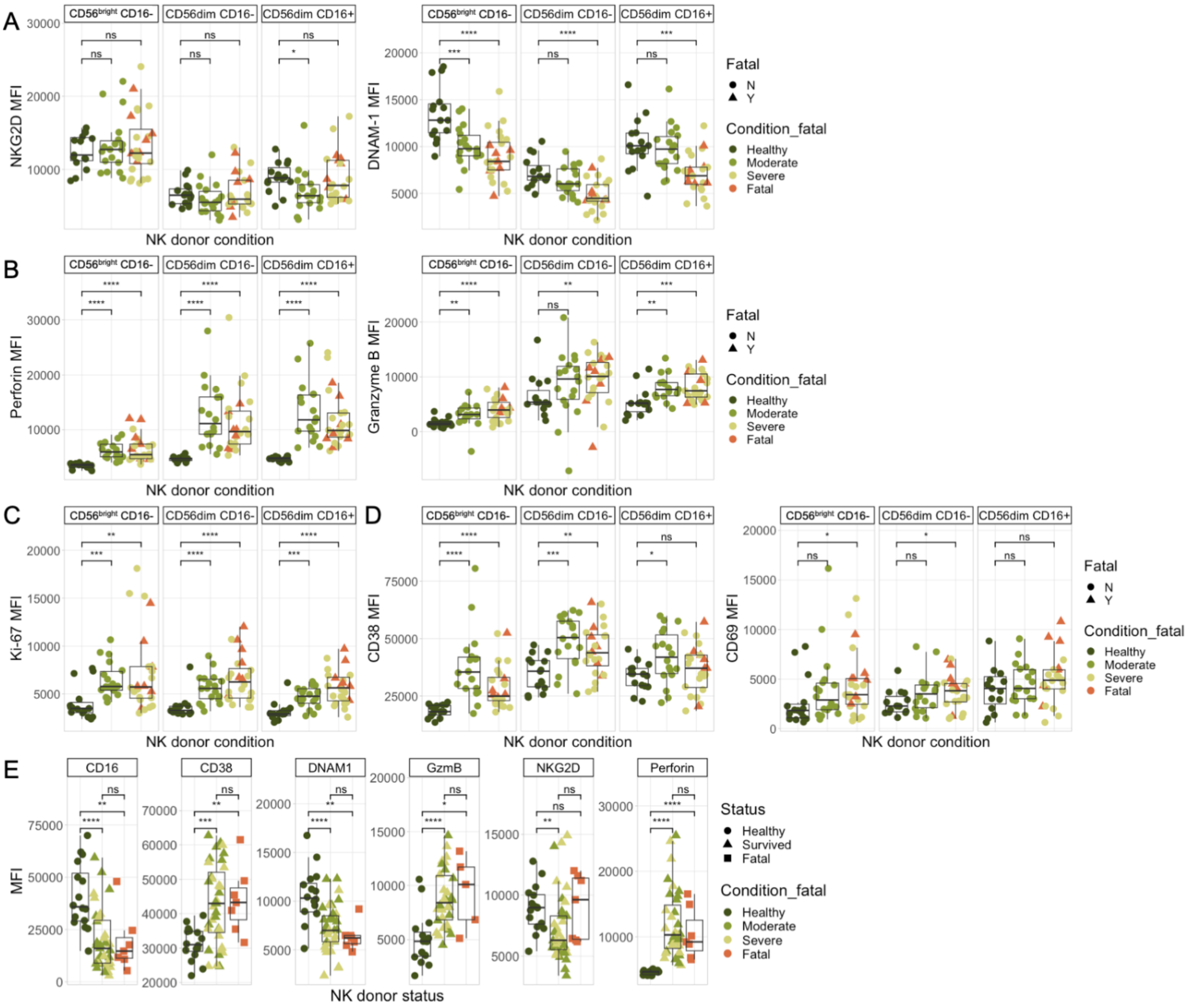


**Supplementary Figure 2: Marker expression in NK cell subsets and comparisons between fatal and non-fatal cases.** A-D) Boxplots showing the MFIs of each marker shown in main text Fig. 2 across three major NK cell subsets ($CD56^{+}CD16^{-}$, left; $CD56^{dim}CD16^{-}$, center; $CD56^{dim}CD16^{+}$, right) from healthy or COVID-19+ donors. E) Boxplots showing MFIs for each marker in all grouped by survival status. “Healthy” status indicates a healthy control donor; “Survived” status indicates a donor who was hospitalized with COVID-19 and recovered; “Fatal” status indicates a donor who was hospitalized with COVID-19 and did not survive. Statistical significance values for all plots were determined by Wilcoxon rank-sum test.

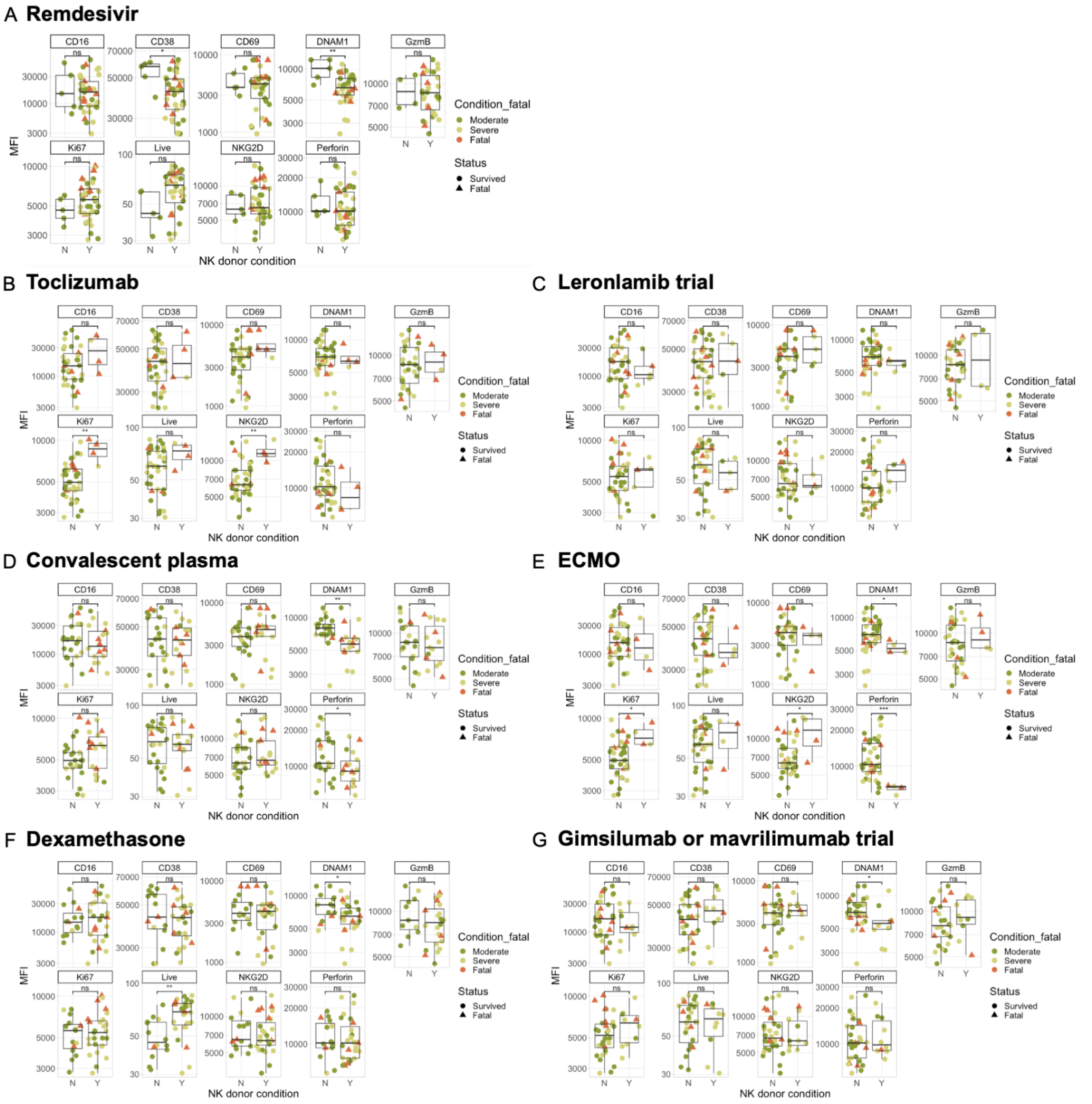


**Supplementary Figure 3: NK cell phenotype by donor treatment status.** A-G) Boxplots showing MFIs of all markers in NK cells in COVID-19 patients who were treated ("Y") or not treated ("N") with each therapeutic used in the cohort. Statistical significance values for all plots were determined by Wilcoxon rank-sum test.

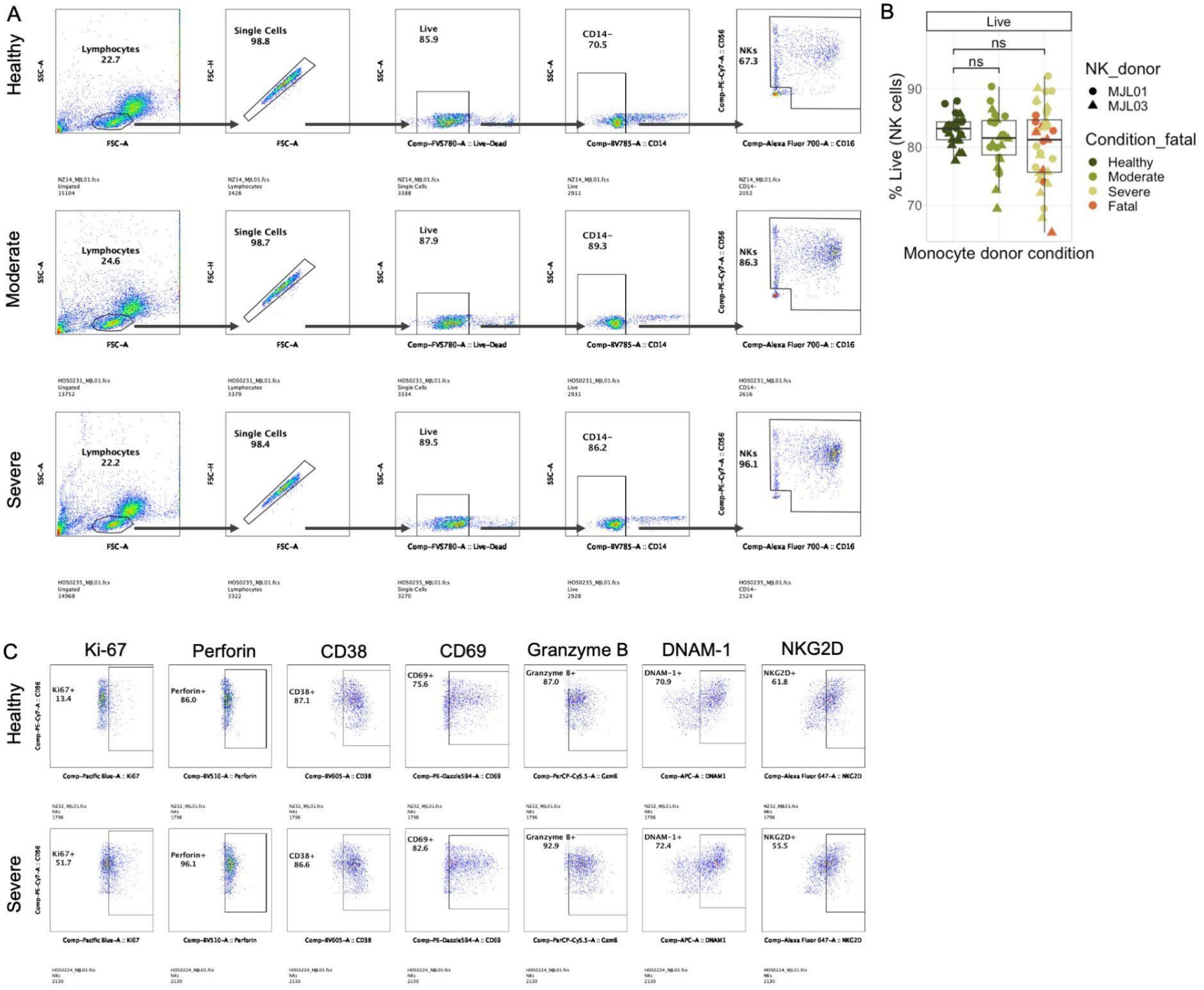


**Supplementary Figure 4: Gating strategy and representative flow plots for direct NK cell-monocyte co-culture experiments.** A) Representative flow plots showing the gating strategy used to identify NK cells in direct NK cell-monocyte co-culture experiments. Example plots are shown for a healthy donor (top), moderate COVID-19 donor (middle), and severe COVID-19 donor (bottom). B) Boxplot showing the percentage of NK cells that were live (negative for eFluor 780 fixable viability dye) in each sample. Statistical significance values were determined using a Wilcoxon rank sum test. C) Representative flow plots showing the expression of all phenotypic markers quantified in main text Fig. 4. Example plots are shown for a healthy donor (top) and severe COVID-19 donor (bottom).

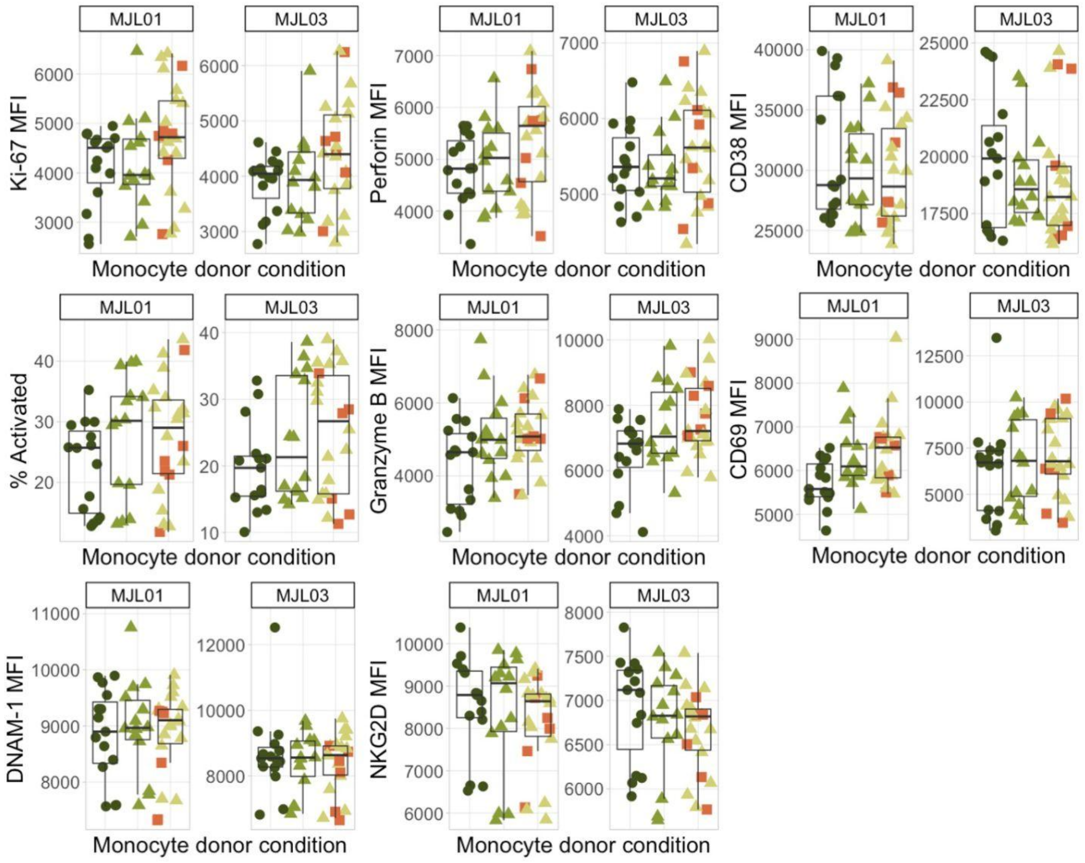


**Supplementary Figure 5: Expression of all markers in the two individual healthy NK cell donors used in direct allogeneic co-culture experiments.** Boxplots showing expression of all markers in the two healthy NK cell donors used for all co-culture experiments following 2-hour co-culture with monocytes from healthy or COVID-19+ donors.

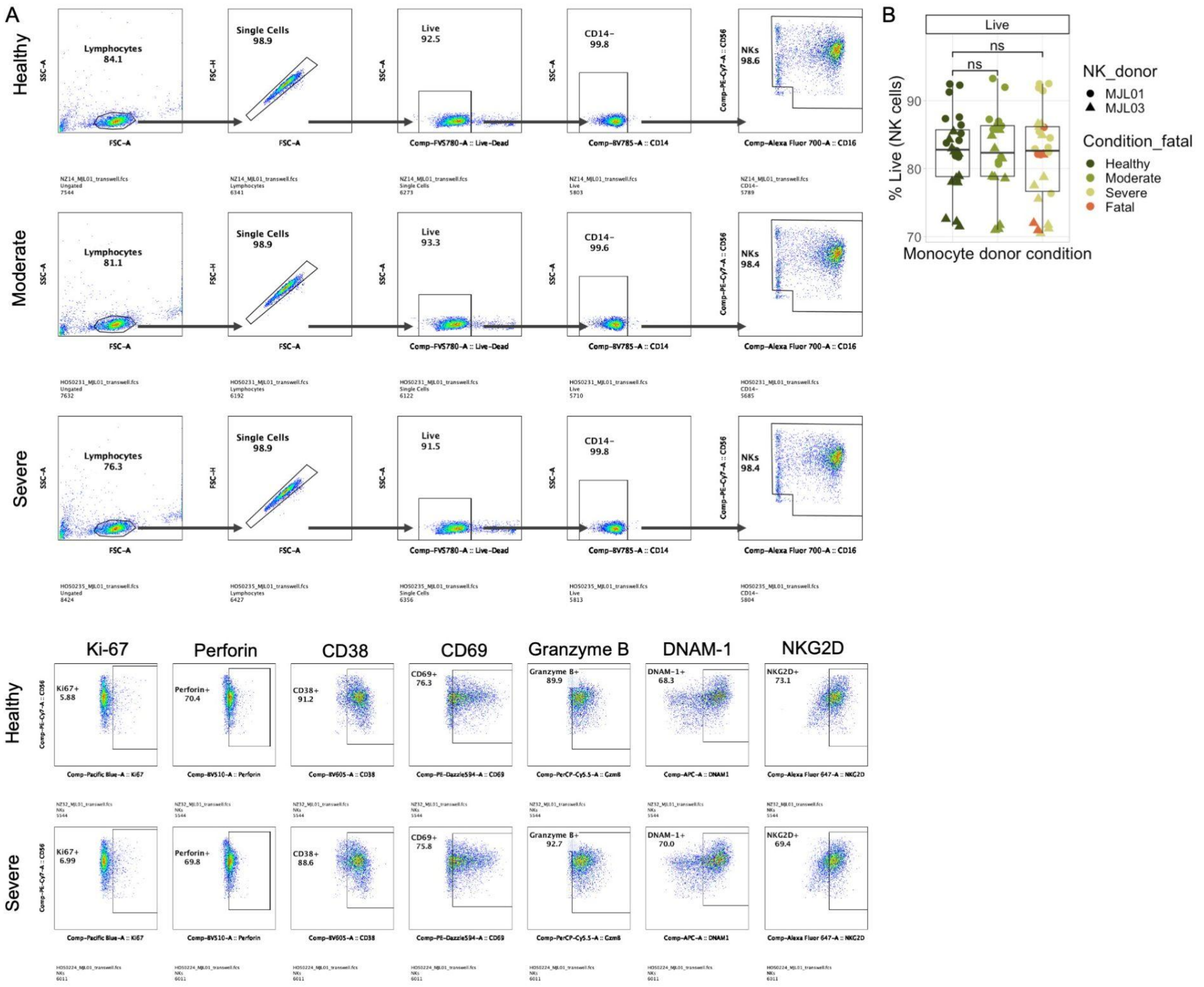


**Supplementary Figure 6: Gating strategy and representative flow plots for transwell NK cell-monocyte co-culture experiments.** A) Representative flow plots showing the gating strategy used to identify NK cells in transwell NK cell-monocyte co-culture experiments. Example plots are shown for a healthy donor (top), moderate COVID-19 donor (middle), and severe COVID-19 donor (bottom). B) Boxplot showing the percentage of NK cells that were live (negative for eFluor 780 fixable viability dye) in each sample. Statistical significance values were determined using a Wilcoxon rank sum test. C) Representative flow plots showing the expression of all phenotypic markers quantified in main text Fig. 5. Example plots are shown for a healthy donor (top) and severe COVID-19 donor (bottom).

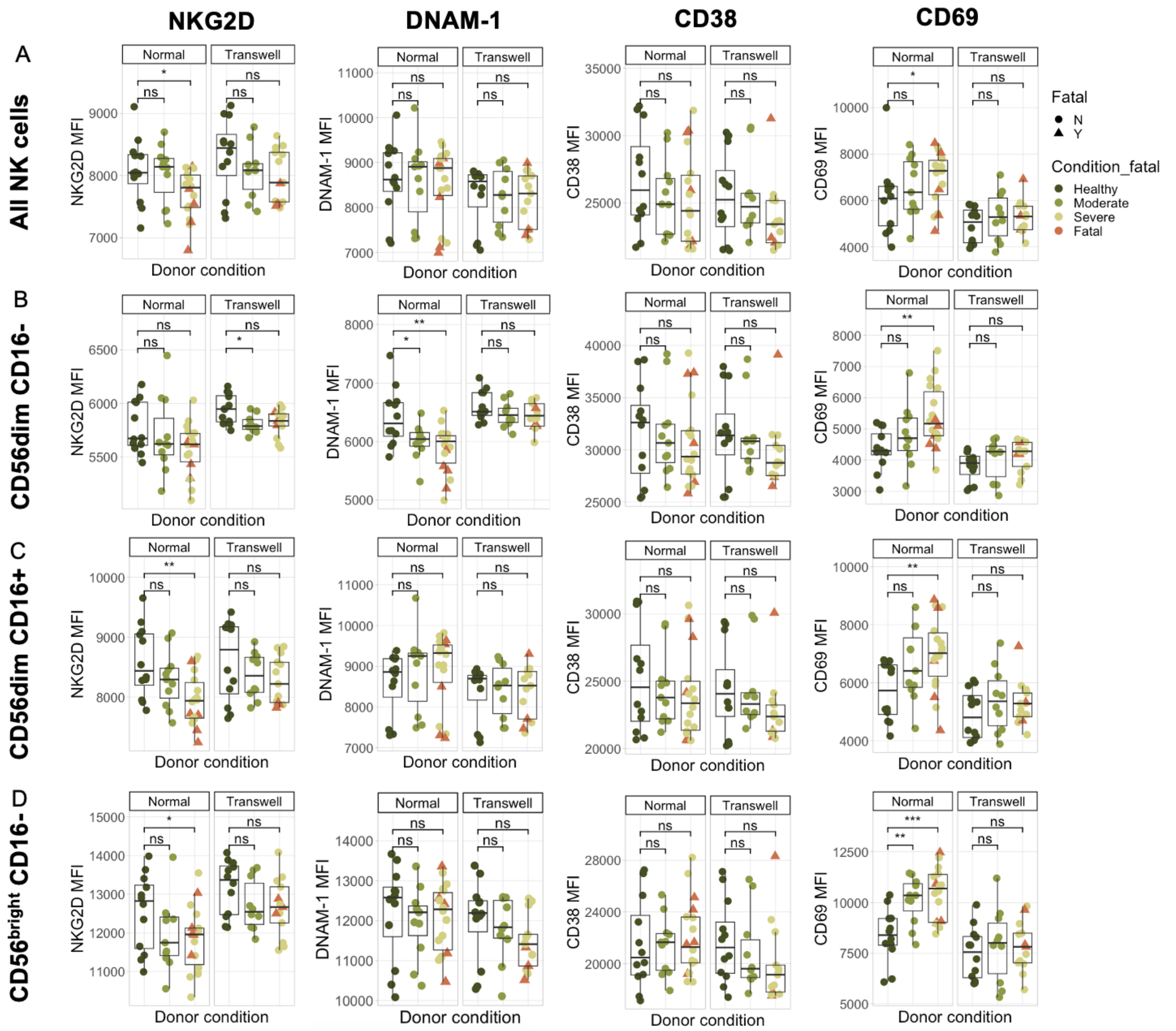


**Supplementary Figure 7: Boxplots showing expression of NKG2D, DNAM-1, CD38, and CD69 in transwell vs direct co-cultures.** A-D) Boxplots showing the MFI of NKG2D, DNAM-1, CD38, and CD69 across severity conditions in normal (round-bottom 96 well plate) cultures and transwell cultures. Each row shows the expression of these markers in a different population of NK cells: total NK cells (A); CD56bright CD16- NK cells (B); CD56dim CD16- NK cells (C); or CD56dim CD16+ NK cells (D). Statistical significance values were determined using a Wilcoxon rank sum test with the Bonferroni correction for multiple hypothesis testing.

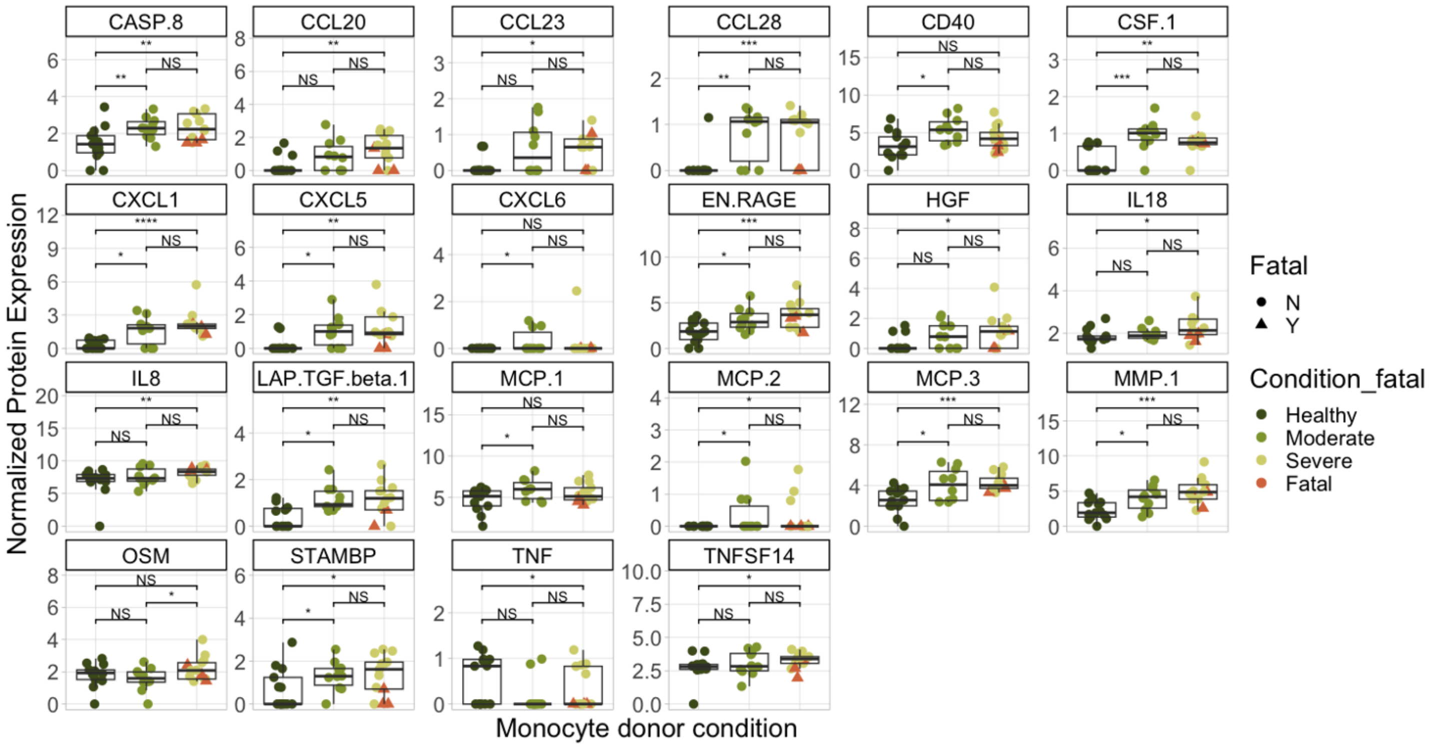


**Supplementary Figure 8: Boxplots showing all O-link analytes whose expressions were significantly different between severity conditions.** Boxplots showing the normalized protein expression of various analytes in the O-link Inflammation Panel across monocyte donor severity conditions. Statistical significance values were determined using a Wilcoxon rank sum test with the Bonferroni correction for multiple hypothesis testing.
